# Cold-Inducible RNA-binding protein (CIRBP) adjusts clock gene expression and REM sleep recovery following sleep deprivation

**DOI:** 10.1101/476911

**Authors:** Marieke MB Hoekstra, Yann Emmenegger, Paul Franken

**Affiliations:** enter for Integrative Genomics, University of Lausanne, Lausanne, Switzerland

**Keywords:** mice, sleep, *Cirbp*, cortical temperature, clock genes, locomotor activity, REM sleep, circadian rhythms

## Abstract

Sleep depriving mice affects clock gene expression, suggesting that these genes partake in sleep homeostasis. The mechanisms linking wakefulness to clock gene expression are, however, not well understood. We propose CIRBP because its rhythmic expression is i) sleep-wake driven and ii) necessary for high-amplitude clock gene expression *in vitro*. We therefore expect *Cirbp* knock-out (KO) mice to exhibit attenuated sleep-deprivation (SD) induced changes in clock gene expression, and consequently to differ in their sleep homeostatic regulation. Lack of CIRBP indeed blunted the SD-incurred changes in cortical expression of the clock gene *Rev-erbα* whereas it amplified the changes in *Per2* and *Clock*. Concerning sleep homeostasis, KO mice accrued only half the extra REM sleep wild-type (WT) littermates obtained during recovery. Unexpectedly, KO mice were more active during lights-off which was accompanied by an acceleration of theta oscillations. Thus, CIRBP adjusts cortical clock gene expression after SD and expedites REM sleep recovery.

## Introduction

The sleep-wake distribution is coordinated by the interaction of a circadian and a sleep homeostatic process (Daan et al., 1984). The molecular basis of the circadian process consists of clock genes that interact through transcriptional/translational feedback loops. CLOCK/NPAS2:BMAL1 heterodimers drive the transcription of many target genes, among which *Period* (*Per1-2*), *Cryptochome* (*Cry1, −2*), and *Rev-Erb* (*Nr1d1, −2)*. Subsequently, PER:CRY complexes inhibit CLOCK/NPAS2:BMAL1 transcriptional activity and thus prevent their own transcription. Through another loop, other clock components such as the transcriptional repressor REV-ERBα regulate the transcription of *Bmal1* (*Arntl*), ensuring together with other transcriptional feedback loops a period of ca. 24 hours (Lowrey and Takahashi, 2011).

The sleep homeostatic process keeps track of time spent awake and time spent asleep, during which sleep pressure is increasing and decreasing, respectively. The mechanisms underlying this process are to date unknown. However, accumulating evidence implicates clock genes in sleep homeostasis [reviewed in (Franken, 2013)]. This is supported by studies in several species (*i.e.* mice, fruit flies and humans), showing that mutations in circadian clock genes are associated with an altered sleep homeostatic response to sleep deprivation (SD) [*e.g.* (Mang et al., 2016, Shaw et al., 2002, Viola et al., 2007, Wisor et al., 2002)]. Furthermore, SD affects the expression of clock genes such as *Rev-erbα, Per1-3* and *Dbp* (Mongrain et al., 2010), but the mechanisms through which this occurs are unclear.

In this study, we examined one such mechanism and hypothesized that some of the SD-induced changes in clock gene expression occur through Cold-Inducible RNA Binding Protein (CIRBP). Decreasing temperature *in vitro* increases CIRBP levels (Nishiyama et al., 1997) and the daily changes in body temperature of the mouse are sufficient to drive robust cyclic levels of *Cirbp* and CIRBP (Morf et al., 2012) in anti-phase with temperature. Although the daily changes in cortical temperature (Tcx) appear circadian, more than 80% of its variance is explained by the sleep-wake distribution in the rat (Franken et al., 1992). Hence, the daily rhythms of cortical *Cirbp* become strongly attenuated when controlling for these sleep-wake driven changes in Tcx by SDs (see Figure 1, based on Gene Expression Omnibus number GSE9442 from Maret et al., 2007). Furthermore, *Cirbp* is the top down-regulated gene after SD (Mongrain et al., 2010, Wang et al., 2010) underscoring again its sleep-wake dependent expression. But how does CIRBP relate to clock gene expression?

**Figure 1.**
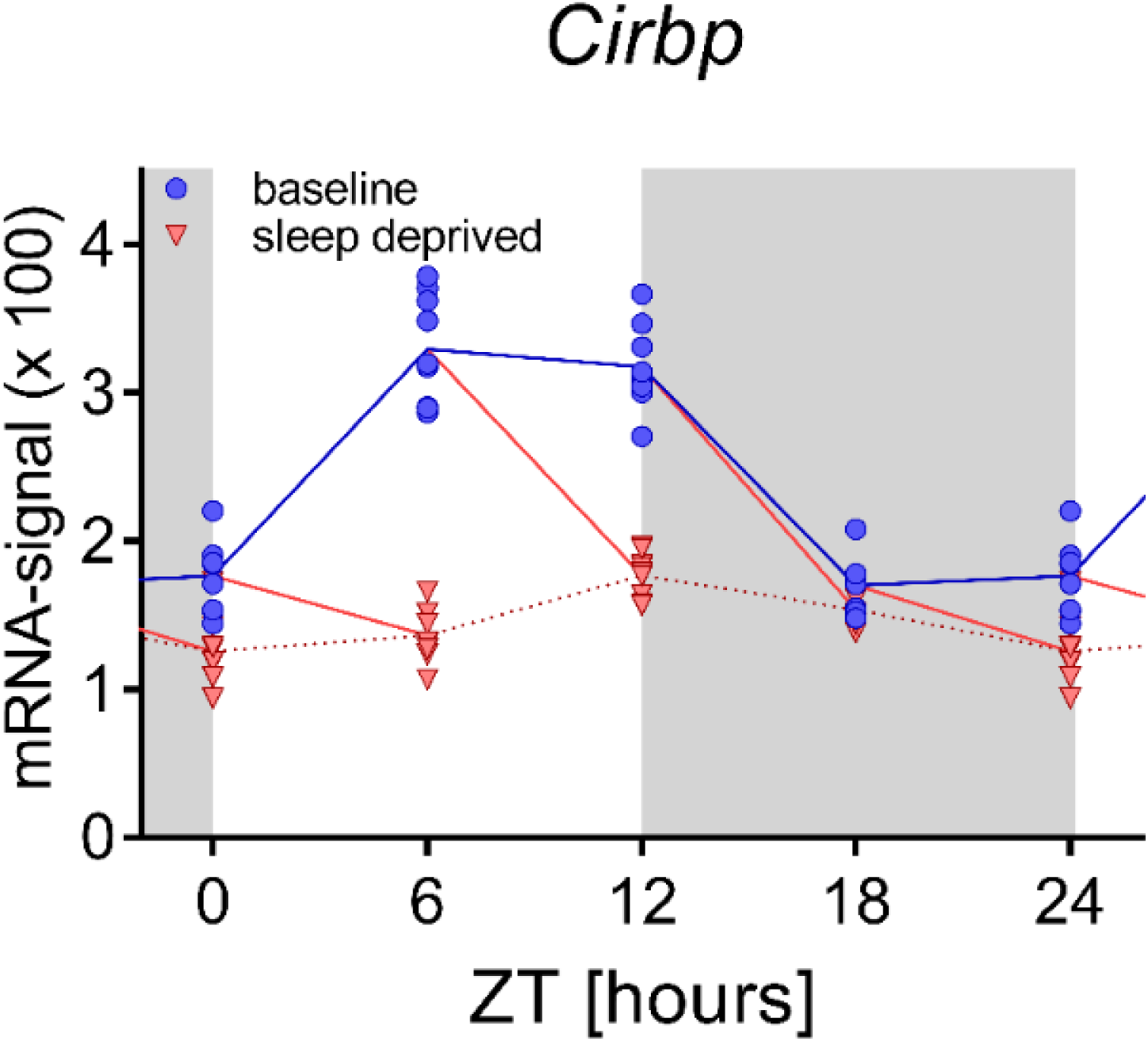
The sleep-wake distribution drives daily changes of central *Cirbp* expression in the mouse. After the onset of the baseline rest phase (ZT0), when mice spend more time asleep and thus T_cx_ decreases, *Cirbp* expression increases (blue symbols and lines), whereas between ZT12-18, when mice spent most of their time awake and T_cx_ is high, *Cirbp* decreases. When controlling for these diurnal changes in sleep-wake distribution by performing four 6h SD [sleep deprivation] starting at either ZT0, −6, −12, or −18, the diurnal amplitude of *Cirbp* is greatly reduced (red symbols represent levels of expression reached at the end of the SD). Nine biological replicates per time point and condition from three different inbred strains of mice were used (one data point missing at ZT18), and RNA was extracted from whole brain tissue (see Maret et al. 2007 for details). Data were taken from GEO GSE9442 and accessible in source file 1.

Two independent studies showed that the temperature-driven changes in CIRBP are required for high amplitude clock gene expression in temperature synchronized cells (Morf et al., 2012, Liu et al., 2013). Therefore, we and others (Archer et al., 2014) hypothesized that changes in clock gene expression during SD are a consequence of the sleep-wake driven changes in CIRBP. We used mice lacking CIRBP (*Cirbp* KO) (Masuda et al., 2012) to test this hypothesis. We first assessed whether also in the mouse the daily changes in Tcx are driven by the sleep-wake distribution and what the contribution of locomotor activity (LMA) to these changes was. Next, we assessed SD-induced changes in clock gene expression in WT and KO mice. Because we expected that the response to SD in terms of clock gene expression differed in KO mice, and clock genes partake in sleep homeostasis (Franken, 2013), we also assessed the homeostatic regulation of sleep in *Cirbp* KO and WT mice.

Our experiments revealed that also in the mouse the sleep-wake distribution is the major determinant of changes in T_cx_, with a significant albeit small contribution of LMA. In line with our predictions, we found that lack of CIRBP indeed attenuated the SD-induced changes in the cortical expression of *Rev-erbα* and the homeostatic response in REM sleep time. However, in contrast to our hypothesis, we observed that the changes in *Per2* and *Clock* expression after SD were augmented in *Cirbp* KO mice. Unexpectedly, we discovered that *Cirbp* KO mice were substantially more active compared to their WT littermates without increasing their time spent awake. This dark-phase increase in LMA was accompanied by an acceleration of EEG theta oscillations during active waking. Altogether, our data show that *Cirbp* contributes to some of the SD-induced changes in clock gene expression, but also points to the existence of other sleep-wake driven pathways conveying sleep-wake state to clock gene expression.

## Results

### THE RELATION BETWEEN CORTICAL TEMPERATURE (TCX), SLEEP-WAKE DISTRIBUTION, AND LOCOMOTOR ACTIVITY (LMA)

The dependence of brain or cortical temperature on sleep-wake state has been demonstrated in a number of mammals (Alfoldi et al., 1990, Baker and Hayward, 1968, Deboer et al., 1994, Franken et al., 1992, Hayward and Baker, 1968) but has not been specifically addressed in the mouse. Moreover, no study so far specifically controlled for LMA when quantifying the contribution of sleep-wake state to brain temperature. We therefore measured cortical temperature (T_cx_), LMA and sleep-wake state in WT and *Cirbp* KO mice during two baseline days, a 6hr SD and the following two recovery days. Because the relationship between T_cx_, LMA, and waking in WT and KO mice was alike, we illustrated the results in WT mice only.

#### Fast changes in T_*cx*_ occur at sleep-wake state transitions

A representative example of a 96h recording of LMA, sleep-wake state and T_cx_ is depicted in Figure 2. Consistent with mice being nocturnal animals, the mouse shows more waking and LMA, and overall higher T_cx_ levels during the dark phase. SD on the third recording day (between Zeitgeber Time (ZT)0-6) led to an almost uninterrupted period of 6hr waking, during which LMA and T_cx_ reached values comparable to bouts of spontaneous wakefulness under undisturbed baseline conditions (*i.e*., ZT12-18). A closer inspection of the rapid changes in T_cx_ suggests that sleep-wake state transitions underlie these fluctuations. We further quantified these sleep-wake evoked changes in T_cx_ by selecting and aligning transitions between consolidated bouts of NREM and REM sleep and wakefulness during the two baseline days (Figure 2-B). When entering NREM sleep, T_cx_ consistently decreased, whereas at a transition into wake and REM sleep, T_cx_ increased. The latter transition was characterized by a fast and consistent change in T_cx_; within 1.5 minutes, T_cx_ increased by 0.4°C. The subsequent transition from REM sleep into wake leads to an initial decrease in T_cx_ and contrasts with the waking-evoked increase in T_cx_ when transitioning from NREM sleep to wake. Altogether, these results provide evidence that sleep-wake state importantly contributes to changes in T_cx_. The sleep-wake state evoked changes in T_cx_ did not differ between genotypes (2-way RM ANOVA, factors genotype (GT) and Time; GT: p>0.13, GTxTime: p>0.09).

**Figure 2.**
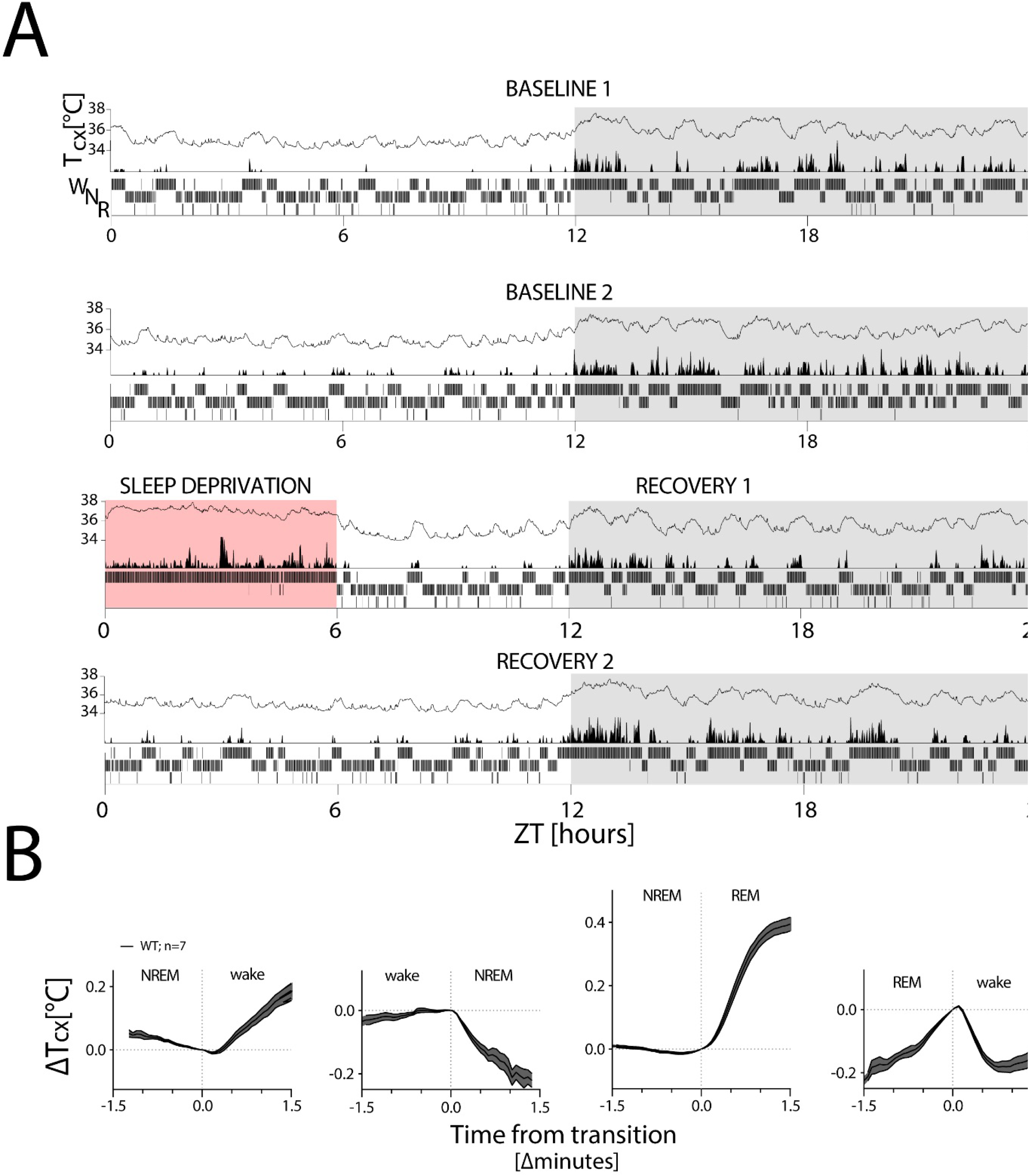
T_cx_ changes with sleep-wake state. T_cx_: cortical temperature; LMA: locomotor activity. (**A)** A representative four-day recording of one mouse in LD 12:12 (in white:grey) during two baseline days (top 2 panels), followed by a 6hr SD (in red; third panel) and two recovery days (bottom 2 panels), with within each panel T_cx_ (top; line graph), LMA (middle; area plot) and sleep-wake states (bottom; hypnogram). Sleep-wake states are averaged per minute to aid visualization. LMA was collected and plotted per minute (see Methods). (**B)** changes in T_cx_, depicted as mean±SEM, relative to T_cx_ at the sleep-wake transition (average of the last value before and first value after transition). T_cx_ increased when transitioning from NREM sleep to wake and to REM sleep (1-way RM ANOVA, factor Time, F(36,180)=40.8, F(44,264)=222; p<0.0001, respectively) and decreased when transitioning from wake to NREM sleep (F(22,132)=1.8, p=0.02). Also the transition from REM sleep to wake affected the time course of T_cx_ (F(23,138)=48.9, p<0.0001). Transition data were obtained from both baseline recordings (see Methods for details).

#### Daily cycles in Tcx are determined by sleep-wake state

After having established that the rapid changes in T_cx_ are indeed evoked by changes in sleep-wake state, we next wondered if the large daily change in T_cx_ is also due to the daily rhythms in sleep-wake state and LMA, and therefore inspected these variables per hour. The LMA data were log2 transformed to allow for parametric assessment. T_cx_, waking and LMA oscillated over the course of the 24h baseline (BL) in similar fashion in both genotypes (2-way RM ANOVA on BL1 and −2 averaged 1h intervals, Factor Time: F(23,207)=70.5; 27.2; 22.5; p<0.0001, respectively, Figure 3-A, Factor GT × Time (1,24); T_cx_: F(23,207)=0.9, p=0.63; waking: 1.7, p=0.03, LMA: 1.21, p=0.24) and the amplitude of the BL change in Tcx did not differ between genotypes (WT: 2.34±0.1, KO: 2.33±0.1; t-test: t(9)=-0.02, p=0.98; average of the difference between the highest and lowest value hourly value in BL1 and −2). Importantly, the time course of waking and LMA both closely resembled that of T_cx_. This observation was supported by the strong correlation between T_cx_ and waking (Figure 3-B left: WT: R^2^=0.76; KO: R^2^=0.81, p<0.0001) and between T_cx_ and LMA (Figure 3-B right: WT: R^2^=0.60; KO: R^2^=0.72, p<0.0001).

**Figure 3:**
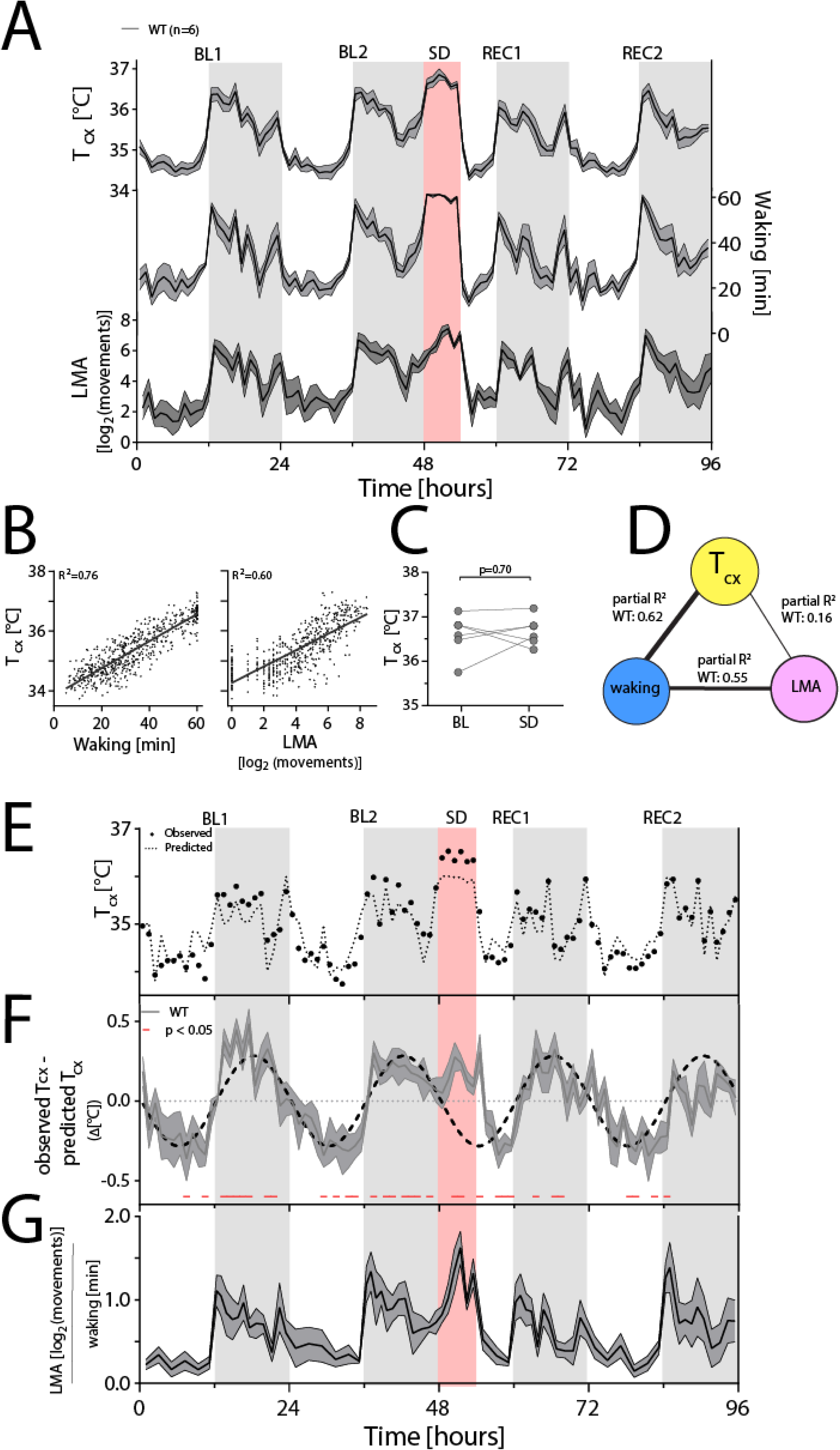
Waking is the major determinant of T_cx_. For panels A, F and G, the dark line represents the mean, areas span ±1SEM. BL: baseline, SD: sleep deprivation, REC: recovery, T_cx_: cortical temperature, LMA: locomotor activity. (**A)** Time course of hourly values of T_cx_, waking and LMA across the entire experiment. (**B)** Both waking (left) and LMA (right panel) correlate with T_cx_ (n=6; 96 values per mouse; R^2^=0.76 and 0.60, respectively, p<0.0001). (**C)** T_cx_ during SD did not differ from levels reached after long waking bouts during BL (t(5)=0.41, p=0.70). (**D)** Waking after correcting for LMA is the major determinant of T_cx_, as revealed by partial correlation analysis; here performed on the combined hourly values of all WT mice. (**E)** a representative example [mouse TC03], with measured T_cx_ (closed circles), and predicted T_cx_ (stippled line) [based on the correlation between T_cx_ and waking]. (**F)** During the dark phase and SD, the expected increase in T_cx_ (based on the amount of waking) is lower than the measured T_cx_, resulting in positive residuals [residuals: observed T_cx_ – predicted T_cx_], whereas during the light phase, the expected T_cx_ is higher than the measured T_cx_, resulting in negative residuals (t-test: data<>0, p<0.05). The sinewave was fitted to the residuals during BL1-2. (**G)** LMA per unit of waking follows a similar pattern as the residuals in figure F.

To assess the influence of waking on T_cx_ at a time of day when T_cx_ is normally low and mice spend most of their time asleep, mice were sleep deprived between ZT0 and ZT6. Mice were 98% of the 6hr SD awake, and thus more awake and active compared to the same time under BL conditions (paired t-test: waking [hours] BL: 2.2±0.1, SD: 5.9±0.04; t(10)=-38.1, p<0.0001; log_2_ [movements], BL: 13.1±2.4, SD: 39.4±2.1; t(10)=-15.2, p<0.0001). These changes led to sustained elevated T_cx_ (average ZT0-ZT6 [°C]: BL: 34.7±0.07, SD: 36.6±0.06, t(10)=-44.3, p<0.0001), suggesting that wakefulness and/or LMA drives changes in T_cx_. Genotype did not contribute to or interact with these changes (2-way ANOVA, GT*SD/BL: p>0.39)

However, factors accompanying the SD other than extended waking, such as stress, could have contributed to the SD-induced changes in T_cx_. To address this issue, we selected within each mouse the longest uninterrupted spontaneous waking bout occurring during BL (average bout length: 100±19 minutes). We then compared T_cx_ during the last 10 minutes of this spontaneous waking bout (to reduce any effects of differences in Tcx at bout-onset) with T_cx_ reached in the last 10 minutes of an equivalent time spent awake from the start of the SD on. T_cx_ reached during the SD and spontaneous wakefulness did not differ (Figure 3-C), also not in KO mice (t(5)=0.84, p=0.44), indicating that factors other than extended wakefulness (*e.g.* light exposure, circadian time, SD-associated stress) do not importantly contribute to the changes in T_cx_ during the SD.

Considering the strong correlation between LMA and T_cx_ (WT: R^2^=0.72; KO: R^2^=0.78; p<0.0001), it could be hypothesized that LMA explains partly the sleep-wake associated changes in T_cx_. To investigate this further, the respective contribution of waking and LMA to changes in T_cx_ was quantified by a partial correlation analysis. Although LMA did significantly contribute, substantially more of the variance in T_cx_ was explained by waking in both genotypes (paired t-test on Fisher Z-transformed R^2^-values from each individual mouse’s partial correlation on hourly waking and T_cx_, and on hourly LMA and T_cx_: WT: t(5)=5.1, p=0.004; KO: t(5)=10.7, p=0.0001; see also Figure 3-D for R^2^-partial correlation coefficients which are based on hourly data from all WT mice combined). We then determined the variance that could not be explained by the correlation between waking to T_cx_ (*i.e.* the residuals) by calculating the difference between the observed T_cx_ in a given hour and the predicted T_cx_ based on the time-spent-awake in that hour. The linear regression overestimated and underestimated T_cx_ during the light phase and dark phase, respectively (Figure 3-E,F; BL1 and BL2), leading to negative residuals during the light phase and positive residuals during the dark phase. Fitting a sinewave through the residuals of the two baseline days revealed a ‘circadian’ distribution, with a mean amplitude of 0.29°C, which is almost twice the amplitude as was previously reported on in the rat [0.15°C] (Franken et al., 1992). Interestingly, when considering the time course of the residuals throughout the experiment, including the SD and recovery, a consistent parallel with the distribution of LMA expressed per unit of waking became evident (Figure 3-G).

Therefore, to determine if including LMA in addition to waking, could predict a larger portion of the variance in T_cx_, we applied three Mixed Linear Models, where LMA was considered by expressing LMA per unit of waking (LMA/Waking). Model1 explained the variance in T_cx_ based on waking alone, Model2 also incorporated LMA/Waking, and Model3 considered additionally the interaction between Waking and LMA/Waking. Indeed, Model3 predicted best the variance in T_cx_ although in terms of explaining the variance in T_cx_, the improvement is marginal over the two other models (Model1: R^2^_c_ =0.84; Model2: R^2^_c_ =0.85; Model3: R^2^_c_ =0.86; chi-squared test: Model1 vs Model2: *X*^2^(5)=16.2; p<0.0001; Model2 vs Model3: *X*^2^(6)=25.0; p<0.0001). Thus, the sleep-wake distribution is the most important determinant of T_cx_ but LMA during waking is modestly contributing as well. Nevertheless, also the residuals of this model, depicted in Figure 3, supplement 1, still showed a similar pattern like the residuals in Figure 3-F, pointing towards the contribution of other (circadian) variables and/or a non-linearity of the association between the contribution of LMA and sleep-wake states to changes in T_cx_.

### THE INFLUENCE OF SD AND CIRBP ON TRANSCRIPTS IN CORTEX AND LIVER

After establishing that also in the mouse the sleep-wake distribution is the major determinant of Tcx, we assessed whether the SD-incurred changes in CIRBP participate in linking the effect of SD to clock gene expression. To this end, we quantified 11 transcripts from liver and 15 from cortex before and after SD by RT-qPCR. Genes of interest included transcripts affected by SD (Maret et al., 2007, Mongrain et al., 2010) and/or by the presence of CIRBP (Liu et al., 2013, Morf et al., 2012), with an emphasis on clock genes. Mice were sacrificed before SD at ZT0, or 6 hours later after SD (ZT6-SD) together with non-sleep deprived control mice that could sleep *ad lib* (ZT6-NSD). Statistics on ZT0 (t-test) and ZT6 (2-way ANOVA) can be found in Table 1.

**Table 1:**
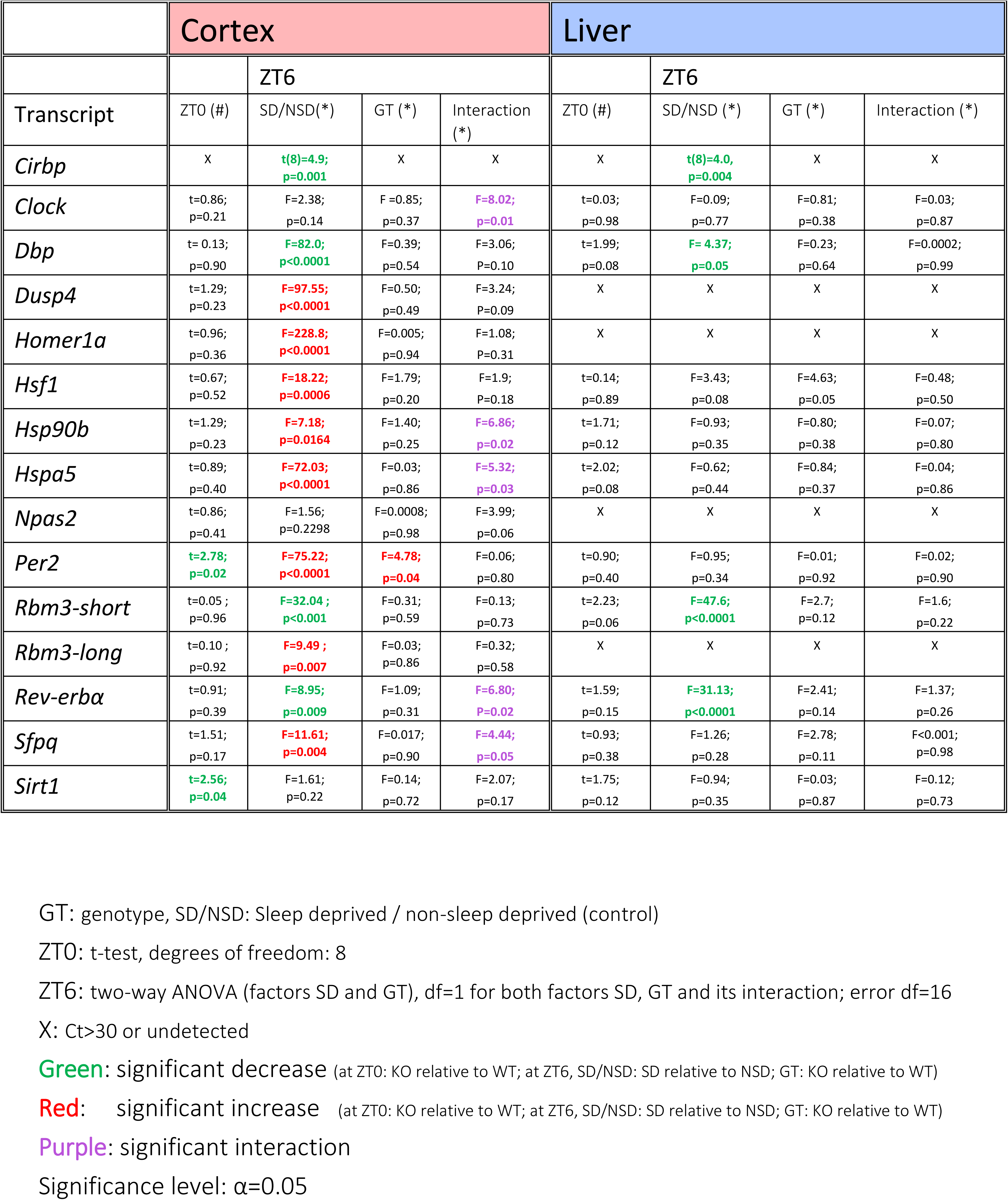
statistics on RT-qPCR results

From ZT0 to ZT6 under undisturbed conditions, T_cx_ decreased because mice spend more time asleep compared to the previous hours in the dark phase (see also Figure 3-A). This decrease in T_cx_ was accompanied by the expected increase of the expression of the cold-induced transcript *Cirbp* in WT mice (cortex: t(8)=3.2, p=0.01; liver: t(8)=2.7, p=0.03; Figure 4-A and Figure 4, supplement 1, compare also with the time course of *Cirbp* Figure 1). In contrast, SD during the same time span incurred a decrease in cortical and hepatic *Cirbp* relative to non-sleep deprived controls (cortex: Figure 4-A; liver: Figure 4, supplement 1), consistent with the wake-induced increase in Tcx during SD. No *Cirbp* mRNA was detected in KO mice.

**Figure 4.**
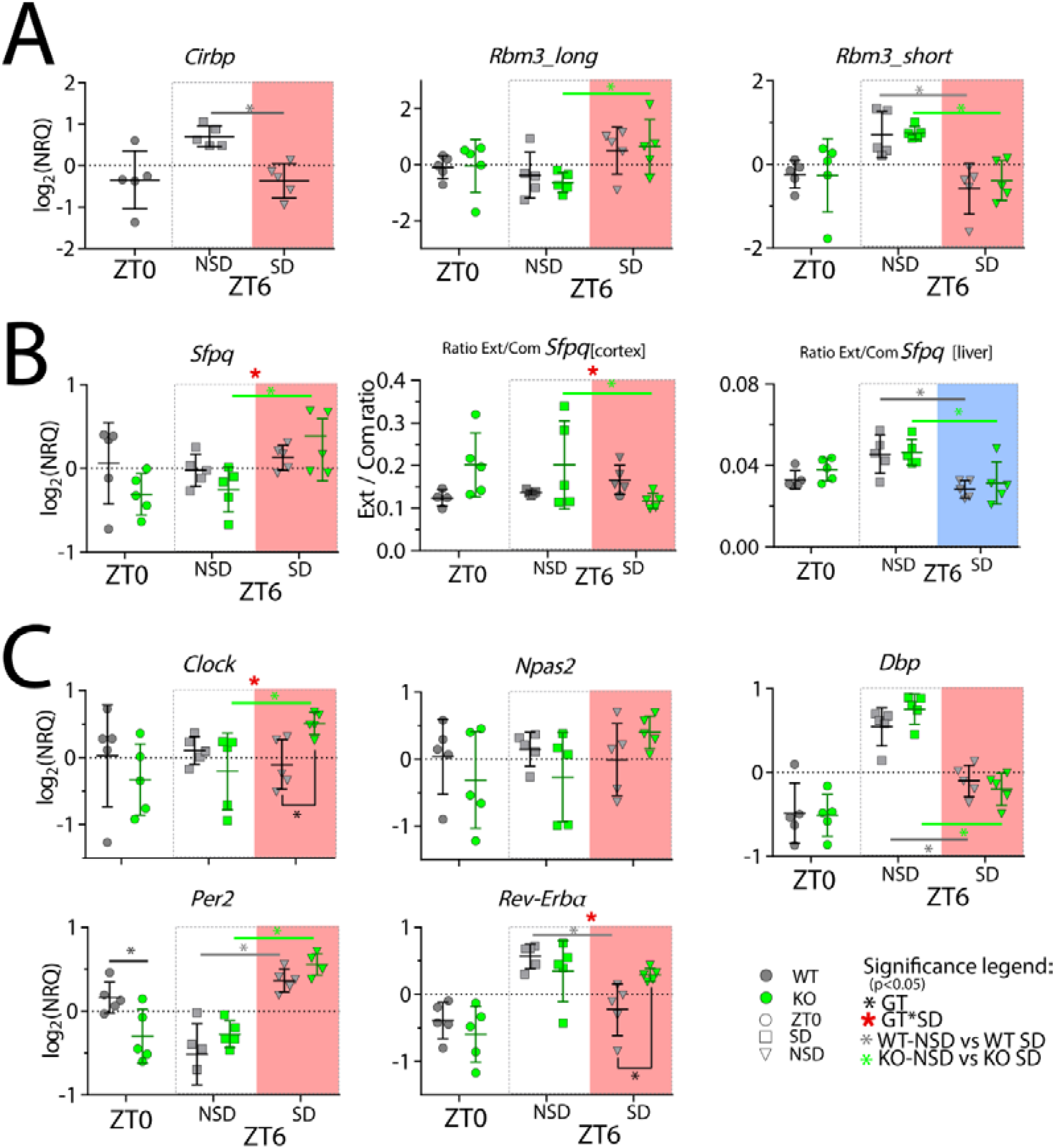
Cortical expression of several genes is affected by SD and the lack of CIRBP. NRQ: Normalized Relative Quantity, SD: sleep deprivation, GT: Genotype. n=5 for each group, each symbol represents an observation in one mouse. Mice were sacrificed at ZT0, at ZT6 after sleep deprivation (ZT6-SD) or after sleeping *ad lib* (ZT6-NSD). Statistics are performed separately for ZT0 (factor GT, t-test), and ZT6 (factor GT and SD; 2-W ANOVA). Significant (p<0.05) GT differences are indicated by a black line and *, the effect of SD in WT mice with a grey line and *, and in KO mice with a green line and *. Interaction effects (GTxSD) at ZT6 are indicated by a red *. See Table 1 for statistics.

RBM3 is another cold-inducible RNA Binding Protein and, like CIRBP, conveys temperature cycles into high-amplitude clock gene expression *in vitro* (Liu et al., 2013). A long and a short isoform of *Rbm3* (*Rbm3-long* and *–short*, resp.) that differ in their 3’UTR length were discovered in the mouse cortex. Although both isoforms are referred to as ‘cold-induced’, they exhibit opposite responses to SD (Wang et al., 2010), with a decrease in the *short* isoform and an increase in the *long* isoform. We found that overall, the short isoform was more common than the long isoform in the cortex (PCR cycle detection number for all samples pooled: cortex: *Rbm3-short*: 25.6±0.2, *Rbm3-long*: 29.7±0.1, amplification efficiency *Rbm3-short*: 2.11 and *Rbm3-long* 2.07). In the liver, only the short isoform was detected (liver: *Rbm3-short*: 28.2±0.2, *Rbm3-long*: >32; *i.e.*, beyond reliable detection limit). We confirmed that after SD, *Rbm3-short* was decreased in the cortex (Figure 4-A) and liver (Figure 4, supplement 1), whereas *Rbm3-long* was increased in cortex. The latter observation reached significance only in the KO mice (Figure 4-A).

As anticipated, cortical expression of the activity (and waking)-induced transcripts *Homer1a, Dusp4, Hspa5/BiP, Hsp90b*, and *Hsf1* was increased by SD (Figure 4, supplement 2). Post-hoc tests revealed that the latter two were significantly increased only in *Cirbp* KO mice. Furthermore, the effect of SD on the transcripts *Hsp90b* and *Hspa5* was significantly amplified in *Cirbp* KO mice compared to WT mice. Unexpectedly, no changes in the expression of heat shock transcripts incurred by SD or genotype were detected in the liver (Figure 4, supplement 1).

*In vitro* studies have shown that the presence of CIRBP is associated with longer 3’UTRs of its target genes, such as the transcript splice-factor proline Q (*Sfpq*), resulting in a higher prevalence of long isoforms (extended or *ext*) over all isoforms (common or *com*), and thus an increased *ext/com* ratio [see FigS4-S5 in (Liu et al., 2013)]. We therefore expected a lower *ext/com* ratio in mice lacking CIRBP. However, under baseline conditions [ZT0 and ZT6-NSD], *Cirbp* KO mice did not differ in their *ext/com* ratio from WT littermates (ZT0: liver: (t(8)=1.55, p=0.16; cortex: t(7)=2.0, p=0.09; ZT6-NSD: liver: t(8)=0.19, p=0.85, cortex: t(8)=1.4, p=0.20). Because also RBM3 partakes in determining the *ext/com* ratio (Liu et al., 2013), the lack of an effect of CIRBP on the *ext/com* ratio could be due to compensation by RBM3. We could test this by assessing the effect of SD on the *ext/com* ratio, because SD acutely suppresses both RBM3 and CIRBP. Indeed, SD significantly decreased the *ext/com* ratio in the liver in both genotypes (Figure 4-B; 2-way ANOVA, factor SD: F(1,16)=20.4, p=0.003). In the cortex, however, we observed an unexpected non-significant increase in WT mice and a significant decrease in KO mice, leading to a significant GT x SD interaction (cortex: F(1,16)=5.25, p=0.036). Therefore, our data are inconclusive in confirming a role for CIRBP, and possibly RBM3, in the *in vivo* determination of *Sfpq*’s *ext/com* ratio.

Our main question concerned the contribution of CIRBP to sleep-wake induced changes in clock gene expression. Previous studies evaluating the effects of SD on cortical clock transcripts showed a consistent increase in *Per2* and a decrease in *Dbp* and *Rev-erbα* whereas the response of *Clock* and *Npas2* varied among studies, but if any, tended to increase after SD (reviewed in (Mang and Franken, 2015)). Indeed, in the cortex of WT mice, SD increased cortical *Per2*, decreased *Dbp* and *Rev-erbα* and did not significantly affect *Clock* and *Npas2* (Figure 4-C). In accordance with our hypothesis, CIRBP attenuated the SD induced changes of cortical *Rev-erbα*, a transcriptional repressor recently implicated in the sleep homeostat (Mang et al., 2016). This observation contrasts with the genotype-dependent changes in *Per2*, because when considering the lower levels of cortical *Per2* in *Cirbp* KO mice at ZT0, the effect of SD was amplified (Figure 4-C, 2-way ANOVA, ZT0-ZT6[SD], interaction effect GT × SD: F(1,16)=12.4, p=0.003). Also, the expression of *Clock* in the cortex was significantly increased by SD in *Cirbp* KO mice and not in WT littermates.

Compared to the cortex, the clock gene expression in the liver appeared more resilient to the effects of SD, as only *Dbp* and *Rev-erbα* were significantly affected and not *Per2* (Figure 4, supplement 1). The lack of CIRBP did not interfere with this response, nor did it contribute to genotype dependent changes of other (clock) gene transcripts in the liver.

Taken together, the absence of CIRBP modulated the SD induced changes in the cortical expression of the clock genes *Rev-erbα, Clock* and *Per2*. Furthermore, the expression of transcripts in the heat shock pathway were also affected in a genotypic manner by SD.

### CIRBP CONTRIBUTES TO SLEEP HOMEOSTASIS

Because *Cirbp* KO mice showed a modulated response to SD in three out of the five cortical clock gene transcripts we quantified, and clock genes importantly partake in the sleep homeostatic process (reviewed in (Franken, 2013)), we hypothesized that *Cirbp* KO mice have a blunted sleep-homeostatic process. We quantified EEG power in the delta band [0.75 – 4.0 Hz] during NREM sleep, which is a proxy of NREM sleep pressure and reflects a homeostatically regulated sleep process, Process S (Daan et al., 1984). As additional sleep homeostatic measures, we calculated the amount of NREM and REM sleep recovered after SD relative to baseline sleep.

#### Baseline characteristics of sleep-wake behavior do not differ between Cirbp KO and WT mice

During the two baseline days, no significant differences in waking, NREM or REM sleep were observed. This was neither the case for time spent in these three behavioral states per light and dark phase (Table 2), nor for the distribution of sleep and waking across the day (see Figure 5-A and Figure 6-A). Noteworthy is that under constant darkness we did not detect a change in circadian period length (period (hours): WT (n=5): 23.8±0.03 and KO (n=7): 23.8±0.01).

**Table 2.**
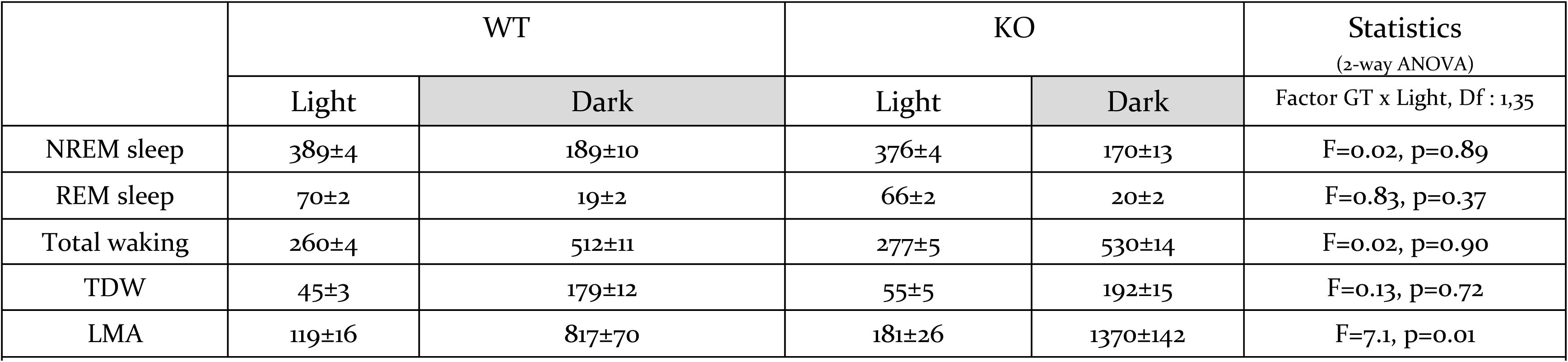
Baseline time spent in sleep-wake states (min) and LMA (movements) per 12 hours per genotype, averages of BL1-2. 2-way ANOVA (Factor GT and Light/Dark) on those same 12-hour values. Degrees of freedom for both GT and Light/Dark: Df=1; error term: Df=35.

**Figure 5.**
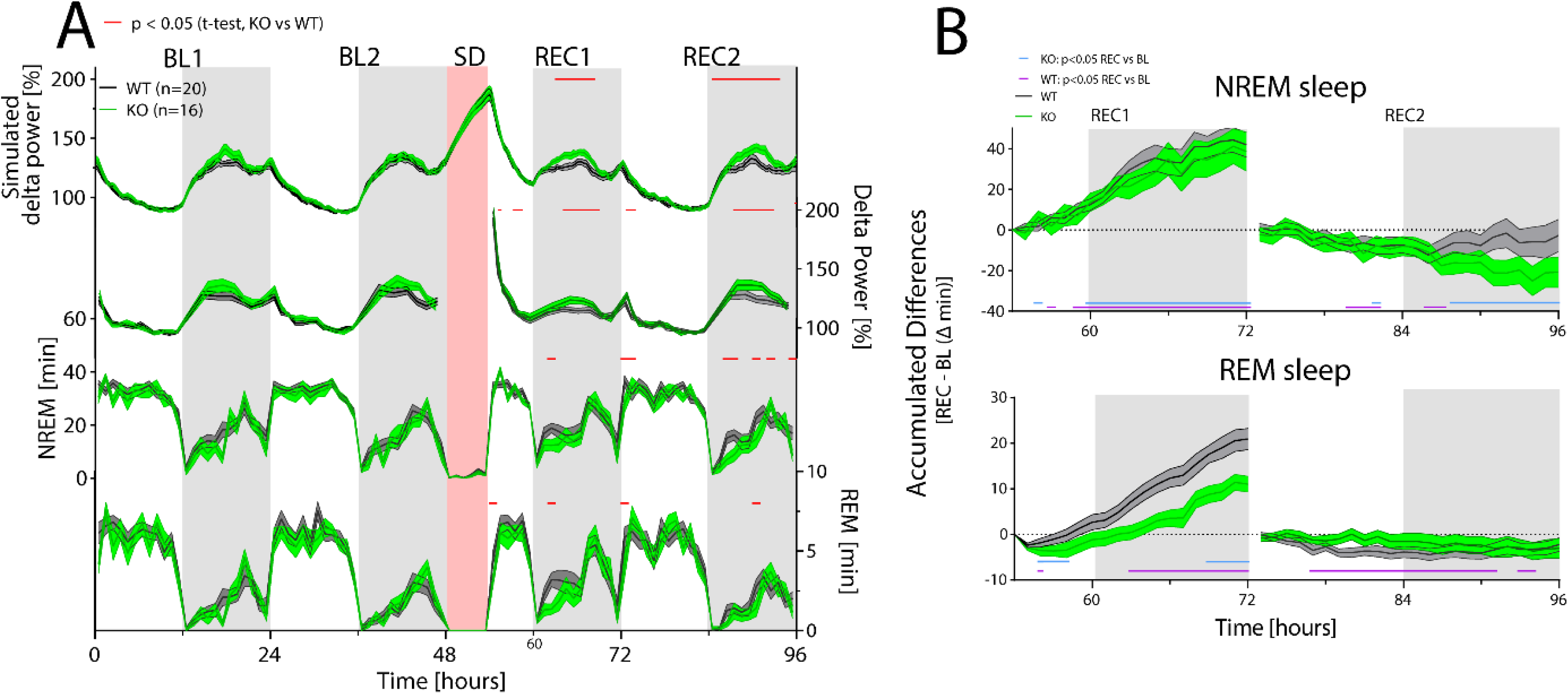
CIRBP modulates the sleep-wake distribution and REM sleep recovery after sleep deprivation. *Cirbp* KO (green lines and areas) and WT (black line, grey areas) mice during the two baseline days (BL1 and −2), sleep deprivation (SD), and the two recovery days (REC1 and −2; areas span ±1SEM range). (**A)** From top to bottom: Simulated delta power (Process S), measured NREM delta power, NREM sleep and REM sleep. During BL, trends, but no significant effect of GT or its interaction with time were detected in the simulation of process S, delta power, NREM sleep or REM sleep. During REC, the simulation predicts increased delta power in *Cirbp* KO mice (GT: F(1,34)=5.56, p=0.024), based on differences in NREM sleep distribution (GT: F(1,34)=6.02, p=0.0194) which are also reflected by increased delta power during the dark phase (GT: F(1,34)=4.65, p=0.038). GT effects in REM sleep were also detected during recovery (factor GT: F(1,34)=5.45, p=0.026). Exact timing of GT differences is indicated by superimposed red lines (post-hoc t-test, p<0.05). (**B)** Top: KO mice recover as much NREM sleep as WT mice in the first 18hrs after SD (REC1 at 72Hr: WT: 41.9±6.1 KO: 38.6±9.7 min; t-test: t(34)=0.30, p=0.76). Bottom: KO mice accumulated less REM sleep during the first recovery day over the baseline day in comparison to WT mice relative to baseline (REC1 at 72Hr, WT: 20.9±2.3 KO: 9.9±2.0, t-test: t(34)=3.7, p=0.0007).

**Figure 6.**
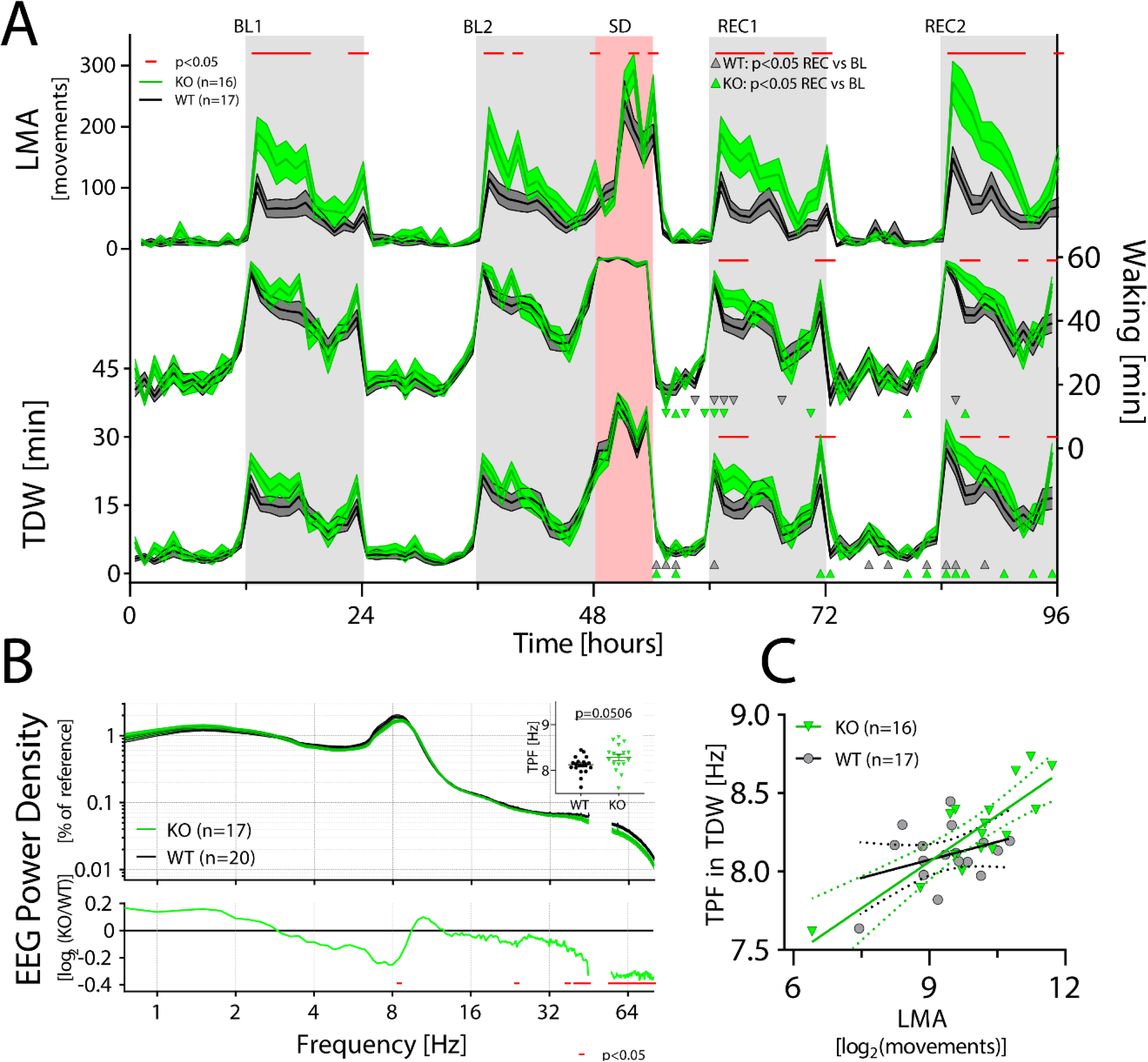
CIRBP suppresses LMA and affects spectral composition during TDW. LMA: locomotor activity, TDW: theta dominated waking; TPF: theta peak frequency. (A) *Cirbp* KO (green lines and areas) and WT (black line, grey areas) mice during the two baseline days (BL1 and −2), sleep deprivation (SD), and the two recovery days (REC1 and −2; areas span ±1SEM range). (**A)** *Cirbp* KO mice are more active in the dark periods only (BL: GTxTime: F (47, 1457) = 3.5, P<0.0001; REC: GTxTime: F (41, 1271) = 5.2, P<0.001), and spent more time awake and in theta dominated waking (TDW) during REC compared to WT mice (total waking: BL: GTxTime: F(47,1457)=1.1, P=0.33 REC: GTxTime: F (41, 1271) = 1.9, P=0.0005; TDW: BL: GTxTime: F (47, 1457) = 1.1, P=0.35; REC: GTxTime: F (41, 1271) = 1.8, P=0.0025). Significant genotype differences are marked superimposed red lines (post-hoc t-tests; p<0.05). Δ and ∇ indicate a significant increase and decrease in REC compared to same time in BL, respectively. (**B)** CIRBP contributes to the spectral composition of TDW in the dark phase (2-way RM ANOVA; GTxFreq: F(278,9730)=2.0; p<0.0001, red symbols in lower panel: post-hoc t-tests, p<0.05), and KO mice tend to have faster TPF during TDW in the dark phase (t(35)=2.0; p=0.0506). (**C)** TPF in the dark phase correlates only in the KO mice significantly with LMA (WT: R^2^=0.12, p=0.17, KO: R^2^=0.71, p<0.0001).

#### Sleep homeostatic processes under baseline and recovery

The time course of delta power in the two genotypes was overall similar. In the dark phase, when mice spent most of their time awake and thus sleep pressure accumulates, delta power during NREM sleep was highest. This contrasts with the end of the light phase [ZT8-12], where NREM sleep delta power reached its lowest levels of the day due to the high and sustained prevalence of NREM sleep in the preceding hours. Despite the overall similarities in daily changes of NREM delta power, subtle differences were observed: delta power levels were higher during the dark phase in *Cirbp* KO compared to WT mice, and these differences reached significance during the dark periods of recovery (Figure 5-A, 2^nd^ graph from top).

Differences in delta power can be attributed to changes in the dynamics of the underlying homeostatic process, Process S, and/or to changes in the sleep-wake distribution. Evidence supporting the latter possibility was observed because *Cirbp* KO mice tended to spend less time in NREM sleep (and more time awake) during the early dark phase compared to WT mice, reaching significance during the recovery (Figure 5-A; 3^rd^ graph from top). To test if these changes in the sleep-wake distribution were indeed sufficient to raise NREM delta power above WT levels, we estimated the increase (τ_*i*_) and decrease (*τ*_*d*_) rate of delta power by a simulation of Process S based on the sleep-wake distribution. We assumed Process S to increase exponentially during waking and REM sleep by time constant τ_*i*_ and to decrease during NREM sleep by time constant *τ*_*d*_ (see Materials and Methods, and (Franken et al., 2001) for more details). This simulation not only captured well the overall dynamics (mean square of the measured-predicted differences, mean±SEM: WT: 10.1±0.3, KO: 10.4±0.4) but also the genotype differences in delta power (Figure 5-A; top graph). No differences in the time constants of Process S were detected (see Table 3). Hence, the reduction in NREM sleep in *Cirbp* KO mice in the beginning of the dark period caused the higher NREM EEG delta power values in subsequent hours, underscoring the notion that small differences in NREM sleep time can have large repercussions on delta power when waking prevails and thus Process S increases (Franken et al., 2001).

**Table 3.** Time constants, asymptotes and S_o_ for Process S do not differ between *Cirbp* WT and KO mice. Mean time constants (±SEM) obtained by the simulation (Process S) with the best fit to the NREM delta power values, where the increase of Process S is simulated by τ_i_, the decrease by τ_d_ and the upper-and lower asymptotes by UA and LA, respectively. No significant genotype differences were observed. See material and methods for detailed description of the simulation.

|  | WT | KO | t-test, df=34 |
| --- | --- | --- | --- |
| $S_0$ [%] | $128.2 \pm 2.4$ | $132.1 \pm 2.6$ | $t=1.10$, $p=0.29$ |
| $\tau_i$ [h] | $13.2 \pm 1.2$ | $12.9 \pm 1.0$ | $t=-0.16$, $p=0.87$ |
| $\tau_d$ [h] | $3.0 \pm 0.2$ | $2.8 \pm 0.2$ | $t=-0.74$, $p=0.46$ |
| LA [%] | $45.1 \pm 1.4$ | $45.1 \pm 1.1$ | $t=-0.02$, $p=0.98$ |
| UA [%] | $288.8 \pm 3.0$ | $296.6 \pm 3.2$ | $t=1.80$, $p=0.09$ |

A different aspect of NREM sleep homeostasis concerns the regulation of time spent in this state. This can be quantified by accumulating relative differences in time spent in NREM sleep from corresponding baseline hours over the recovery period. At the end of the first recovery day, both KO and WT mice had gained ca. 40 minutes of NREM sleep relative to baseline (Figure 5-B, upper panel).

The amount of REM sleep is also homeostatically defended (Franken, 2002). At the end of REC1, both WT and KO mice spent more time in REM sleep compared to corresponding baseline hours. However, this increase in REM sleep was significantly attenuated by 46% in *Cirbp* KO mice (Figure 5-B, lower panel). Because no significant differences were detected during baseline in time spent in REM sleep (see also Table 1), this attenuated rebound in REM sleep resulted from less REM sleep during recovery, specifically in the first hours of the dark phase when the genotypic differences were most prominent (Figure 5-A, lowest graph).

Thus, although CIRBP did not affect the processes underlying NREM sleep intensity and NREM sleep time, it did contribute to REM sleep homeostasis by increasing the amount of REM sleep after SD.

### AN UNANTICIPATED WAKING PHENOTYPE IN CIRBP KO MICE

While quantifying sleep-wake states, we observed that *Cirbp* KO mice were more active than their WT littermates during the dark phase (t(31)=-2.56, p=0.015, see also Table 2). More specifically, *Cirbp* KO mice were almost twice as active in the first 6hrs of the dark phase (movements: WT: 463.8±60.7, KO: 801.8±118.4, t(35)=-2.7, p=0.012; Figure 6-A). Interestingly, this pronounced increase was not associated with a significant increase in time spent awake during BL (per 12 hrs: t(35)=1.2, p=0.24, and see Table 2), and indeed *Cirbp* KO mice were more active per unit of waking (average in the dark phase, LMA [movements/waking(min)], WT: 1.3±0.13, KO: 2.1±0.28; t(35)=-2.7, p=0.01). Note that also T_cx_ was not significantly increased in *Cirbp* KO mice during the dark phase (T_cx_: WT: 35.9±0.1, KO: 36.1±0.1, t-test, t(10)=1.3, p=0.24) despite the increased LMA at this time of the day, again underscoring the minimal contribution of LMA to T_cx_.

Because *Cirbp* KO mice are not more awake (Table 2 and Figure 6), we wondered if their increased LMA is associated to the prevalence of sub-states of waking. Theta-dominated waking (TDW) is a sub-state of waking that correlates with activity, prevails during the dark phase and SD, and is characterized by the presence of EEG theta-activity (Buzsáki, 2006, Vassalli and Franken, 2017). Despite their increased LMA, *Cirbp* KO mice did not spend more time in TDW during the dark phase of the BL (see Table 1, t(31)=-1.22, p=0.23). If not time spent in TDW, does the increased LMA in *Cirbp* KO mice relate to changes in brain activity during dark phase TDW?

Although in both genotypes the TDW EEG showed the characteristic theta activity [6.5-12.0 Hz], subtle differences between genotypes were detected in the spectral composition of the EEG signal. Slow [32-45Hz] and fast [55-80Hz] gamma power were both reduced during TDW in *Cirbp* KO mice (Figure 6-B), and this reduction was observed throughout the experiment (Figure 6, supplement 1; and see time course in Figure 6, supplement 2), indicating that these spectral genotype differences are robust across different light conditions, circadian times and throughout the SD.

In contrast, the spectral composition of the EEG during ‘quiet’ waking (*i.e.* all waking that is not TDW) was remarkable similar between the two genotypes (Figure 6, figure supplement 1), demonstrating that the changes in spectral composition of TDW EEG are not the result of a general effect of CIRBP on the waking EEG.

Moreover, we observed a decrease in slow and a non-significant increase in faster theta activity in the TDW EEG of *Cirbp* KO mice, together hinting at an acceleration of theta peak frequency (TPF; lower panel in Figure 6-B). TPF during TDW in BL was indeed increased in KO mice (+0.15Hz) although our significance threshold was not met (t(35)=2.0, p=0.05o6). No suggestions for accelerated TPF in REM sleep during BL, the other sleep-wake state characterized by distinct theta oscillations in the EEG, were detected (WT: 7.43±0.06; KO: 7.56±0.05, t(35)=1.7, p=0.10). During locomotion, increased LMA correlates well with increased TPF (Jeewajee et al., 2008). In accordance with this observation, mean log2-transformed LMA levels per mouse during the dark phase predicted well the mean TPF observed during TDW at the same time of day (WT and KO combined; R^2^=0.52, p<0.0001), although this relationship remained significant only in KO mice when assessing the two genotypes separately (Figure 6-C). This genotype-dependent association between TPF and LMA was not confirmed by a significant difference in slope between the genotypes (ANCOVA, F(1,29)=3.8, p=0.059).

Because the group correlation did not account for inter-individual differences in LMA levels, we also assessed the correlations between TPF and LMA within individual mice. Moreover, to test if this association depended on the lighting condition, we analyzed this during the dark and light phase separately (*i.e.* 24 values per mouse per lighting condition; see also Table 4). In the dark phases, this correlation was significant in all but one mouse (a KO), and both the slope and the predictive power of this correlation did not significantly differ between genotypes (slope: WT: 0.15±0.01, KO: 0.14±0.01, t(31)=0.37, p=0.72; R^2^: WT: 0.81±0.03; KO: 0.79±0.04, t-test on the Fisher Z-transformed R^2^-values: t(31)=-0.49, p=0.62). In the light phases, this association was weaker (dark vs. light: paired t-test: slope: t(32)=7.8, p<0.0001; Fisher Z-transformed R^2^-values: t(32)=5.9, p<0.0001), but again did not differ between genotypes (WT: 0.07±0.01, KO: 0.09±0.01, t(31)=1.2, p=0.23; R^2^: WT: 0.59±0.04; KO: 0.67±0.05, t-test on the Fisher Z-transformed R^2^-values: t(31)=1.2, p=0.23). However, during the light phases, when LMA and TDW are substantially reduced and estimates of TPF during TDW are less precise, we found more non-significant associations in both genotypes (KO: 3/ 16; WT: 3/ 17 mice). Altogether, these results provide further evidence that LMA contributes to TPF and suggests that CIRBP, through its effects on LMA, reduces TPF.

**Table 4:**
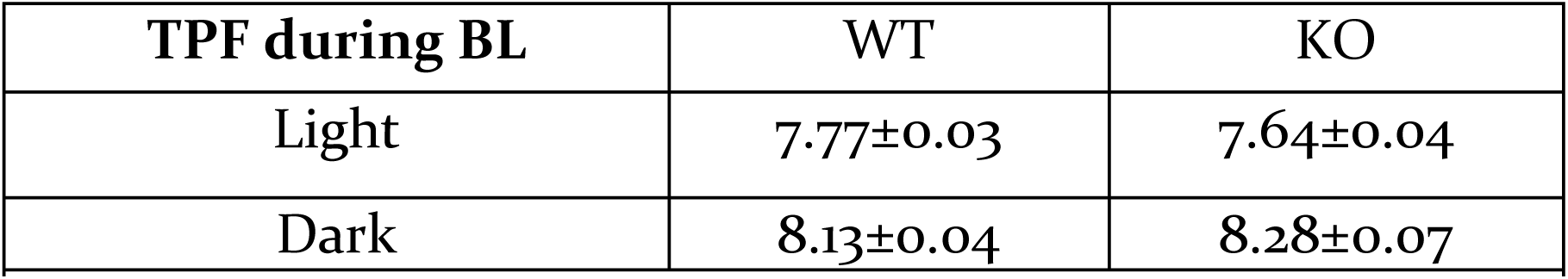
Average TPF during baseline light and dark (mean ± SEM) in TDW

The SD altered the distribution of waking during the recovery relative to BL (3-way RM ANOVA, factor Time, GT and BL/REC, factor BL/REC: REC1: F(1,558)=42.7, p<0.0001; REC2: F(1,1514)=441.8, p<0.0001; see triangles in REC1 and REC2, Figure 6-A). Surprisingly, while time spent awake was overall decreased compared to baseline, we observed several intervals during the recovery in which TDW was increased (3-way RM ANOVA, factor Time, GT and BL/REC, factor BL/REC: REC1: F(1,558)=13.9, p=0.0002; REC2: F(1,1514)=233.8, p<0.0001; Figure 6-A, upwards pointing triangles). This was true for both genotypes. Moreover, genotype differences in the distribution of waking and TDW became significant during the dark phases of both recovery days, with *Cirbp* KO mice spending more time awake and in TDW than WT mice (Figure 6-A; see post-hoc tests indicated by red line), as if SD amplified the non-significant genotype differences during BL (3-way RM ANOVA on hourly values: factor GTxTimexSD: total waking: F(41,1271)=1.4, p=0.04; TDW: F(41,1271)=1.4, p=0.056) but not for LMA (F(41,1271)=1.0, p=0.48).

The EEG spectra during TDW in REC1 and REC2 showed similar profiles as during BL (see Figure 6, figure supplement 2), although there were some changes that in recovery reached significance such as the increase in the delta power band. Along those lines, the non-significant increase in TPF in *Cirbp* KO mice during the BL dark phases became significant in the REC dark phases (REC1: WT: 8.1±0.05, KO: 8.4±0.07, t(35)=2.7, p=0.01; REC2: WT: 8.2±0.05, KO: 8.5±0.08, t(35)=2.6, p=0.01). Also, the non-significant difference in slope at the group level between TPF and LMA during BL (Figure 6-C), became significant after SD (ANCOVA, F(1,29)=5.8, p=0.02), providing further evidence that the suggestive genotype differences under baseline conditions become more pronounced after a challenge of the sleep homeostat.

Taken together, *Cirbp* KO mice were more active during the dark phase, which partly explains the faster TPF. Moreover, KO mice had less EEG power in the gamma band of TDW, and the 6-hour SD strengthened genotype differences in the sleep-wake distribution and EEG activity.

## Discussion

In this study, we showed that, like in other rodents, the sleep-wake distribution is the major determinant of T_cx_ in the mouse. Because of the well-established link between temperature and CIRBP levels, it is likely that the equally well-known sleep-wake driven changes in *Cirbp* expression in the brain are conveyed through the sleep-wake driven changes in brain temperature. As predicted, the SD-incurred changes in the expression of clock genes was modulated by the presence of CIRBP. However, only for *Rev-Erbα* did we observe the anticipated attenuated response to SD in *Cirbp* KO mice, whereas the changes in the expression of *Per2* and *Clock* were amplified compared to WT mice. Moreover, we did discover evidence of altered dynamics of the process regulating time spent in REM sleep. Unexpectedly, *Cirbp* KO mice are more active during the dark phase, and have during TDW reduced power in the gamma band and increased TPF.

### CHANGES IN CORTICAL TEMPERATURE ARE SLEEP-WAKE DRIVEN

When sleep and waking occur at their appropriate circadian times, the changes in both brain and body temperature have a clear 24-hour rhythm and therefore appear as being controlled directly by the circadian clock. However, sleep-wake cycles contribute significantly to both the daily changes in brain and body temperature. In humans, this involvement is powerfully illustrated by spontaneous desynchrony, where body temperature follows both a circadian and an activity-rest (and presumably, sleep-wake) dependent rhythm (Wever, 1979). The contribution of sleep-wake state to the daily dynamics in body temperature is further supported by forced desynchrony studies, such as (Dijk and Czeisler, 1995), estimating that ‘masking’ effects of rest-activity and sleep-wake cycles contributed between 30% and 50% to the amplitude of the circadian body temperature rhythm (Hiddinga et al., 1997, Dijk et al., 2000). Not only in humans but also in smaller animals like rats, a circadian and rest-activity component contribute to the circadian fluctuations in body temperature (Cambras et al., 2007). Thus, the circadian amplitude of body temperature is amplified when wake and sleep occur at the appropriate phase of the circadian rhythm.

In contrast to body temperature, brain temperature in rodents is much more determined by sleep-wake state: 80% of its variance can be explained by the sleep-wake distribution ((Franken et al., 1992) and this study). Likewise, the sleep-wake driven changes in brain temperature are still present in arrhythmic animals (Edgar et al., 1993, Baker et al., 2005), pointing to a more important sleep-wake dependency of brain temperature compared to body temperature. In our study, we also estimated the contribution of LMA to changes in T_cx_ and found that waking with higher LMA is associated with higher T_cx_. Although significant, the contribution of LMA to the daily changes in T_cx_ was modest and explained only 2% more of the variance compared to waking alone. Can we optimize the prediction of T_cx_? A non-linear relationship between sleep-wake state and T_cx_ was assumed previously (Franken et al., 1992) and could have improved the prediction of our model further. This is supported by the residuals from the complete model (see Figure 3, supplement 2), that exhibit under baseline conditions a circadian distribution, whereas during the SD, they remain increased as during the dark phase. Thus, the model overestimates T_cx_ during periods with little waking (light phase) and underestimates T_cx_ during periods that are dominated by waking (dark phase and SD), suggesting a non-linear relationship between these two variables.

Important to consider is that the influence of LMA on T_cx_ is likely affected by the type of activity; for example, rats on a running wheel activity can increase their brain temperature by 2°C within 30 minutes (Fuller et al., 1998). Also exercise in men leads to an increase in (proxies) of brain temperature (Nybo et al., 2002). Thus, although in our study the effect of LMA to T_cx_ was very modest compared to the effect of waking on T_cx_, these contributions likely differ with various types of physical activity.

### LMA-DEPENDENT AND INDEPENDENT CHANGES IN WAKING CHARACTERISTICS

Little is known about the role of CIRBP in neuronal and behavioral functioning. It was therefore unanticipated that *Cirbp* KO mice were more active during the dark phase. Neither were the changes in neuronal oscillations during TDW: a reduction in low- and high gamma power and an increase in TPF. Because increased running speed correlates with increased hippocampal TPF (Jeewajee et al., 2008), and our measured TPF is mainly of hippocampal origin (Buzsáki, 2006), we indeed can relate the increased TPF to the increase in LMA in KO mice. In contrast to TPF, the literature has not consistently reported on a relation between the general decrease in gamma power during active waking and its relation to LMA. Some studies have found that increased speed of movement relates to increased power in the gamma band (Furth et al., 2017, Niell and Stryker, 2010, Vinck et al., 2015), whereas others found that this association is only present in higher gamma frequencies [>60Hz] (Zheng et al., 2015). Thus, it is unclear if LMA relates to changes in gamma power. However, there is a clear increase in power of the high gamma band specifically during the SD (see Figure 6, supplement 2), as noted previously (Vassalli and Franken, 2017). This increase was present in both genotypes suggesting that while KO mice seem to have a reduced capacity to produce fast gamma activity, SD is still able to activate their fast-gamma circuitry. These results, together with the observation that during the light phase the decreased power in the gamma bands was still present at a time of day when LMA did not significantly differ, argue against an association between the decreased power in the gamma bands of *Cirbp* KO mice and their increased LMA.

Interestingly, gamma oscillations are associated with a palette of cognitive processes [reviewed in (Bosman et al., 2014)]. This is further supported by associations between behavioral impairments and changes in gamma power. For example, mice with abnormal interneurons are impaired at the behavioral level (*e.g.* lack of cognitive flexibility) and have a reduction in task-evoked gamma power in their EEG. Pharmacological stimulation of inhibitory GABA-neurons augmented power in the gamma band and rescued the behavioral phenotype of the mutants (Cho et al., 2015).

In the hippocampus, gamma-theta coupling, *i.e.* the occurrence of gamma oscillations at a specific phase of the theta oscillation, has been suggested to aid processes underlying memory [for review see (Colgin, 2015)]. Because CIRBP slows down TPF and increases power in the gamma bands, further analyses and experiments can address if *Cirbp* KO mice have altered phase coherence between these two frequency bands. Together with the postulated function of gamma power in cognitive flexibility, it would be interesting to assess if the spectral phenotype in *Cirbp* KO mice is associated with behavioral abnormalities.

Several aspects of waking that appeared to differ between *Cirbp* KO and WT mice under baseline dark conditions but were non-significant, reached significance during the recovery dark phase. For example, during baseline *Cirbp* KO mice were 4% more awake and 13% more in TDW compared to their WT littermates, which was amplified to 8% and 20%, respectively, during recovery. Also, TPF and the genotype-dependent association between overall TPF and LMA reached significance during the recovery. This suggest that SD amplified the genotypic differences. Other sleep deprivation studies found evidence for similar phenomena, where sleep disturbance can amplify molecular and behavioral phenotypes of Alzheimers’ mouse models (for review, see (Musiek and Holtzman, 2016)) and sensitivity to pain (Sutton and Opp, 2014). Our data indicates that a similar phenomenon occurs in *Cirbp* KO mice, where a single 6-hr SD reveals the suggestive baseline genotypic differences. It would be interesting to understand the dynamics of this change; *e.g.* if they are reversible or if a second SD could augment genotypic differences further.

### CIRBP ADJUSTS CLOCK GENE EXPRESSION AND REM SLEEP RECOVERY FOLLOWING SD

CIRBP modulated the cortical response to SD in the expression of three out of the five clock genes quantified. As anticipated, the SD incurred decrease in cortical *Rev-erbα* was attenuated in *Cirbp* KO mice. REV-ERBα acts as a transcriptional repressor of positive clock elements such as BMAL1 (Preitner et al., 2002). Mice lacking both *Rev-erbα* and its homolog *Rev-erbβ* have a shorter and unstable period under constant conditions and deregulated lipid metabolism (Cho et al., 2012). We recently established that *Rev-erbα* also partakes in several aspects of sleep homeostasis: *Rev-erbα* KO mice accumulate at a slower rate NREM sleep need and have reduced efficiency of REM sleep recovery in the first hours after SD (Mang et al., 2016).

The expression of the clock genes *Per2* and *Clock* was also modulated in the absence of CIRBP, suggesting that parts of the core clock are sensitive to the presence of CIRBP in response to SD. Given the role of clock genes in sleep homeostasis (Franken, 2013), the modulated clock gene expression in KO mice could have contributed to the REM homeostatic sleep phenotype. This is supported by studies showing that mutations in clock genes incurred a loss in REM sleep recovery [*i.e.* CLOCK (Naylor et al., 2000)], or impacted the initial efficiency of REM sleep recovery [*i.e.* DBP (Franken et al., 2000), PER3 (Hasan et al., 2011), and REV-ERBα (Mang et al., 2016). Follow-up studies can address if indeed the changes in clock gene expression in *Cirbp* KO mice are functionally implicated in the REM sleep phenotype.

The other aspects of the homeostatic regulation of sleep that we inspected, NREM EEG delta power and time spent in NREM sleep after sleep deprivation, were unaffected in *Cirbp* KO mice. Thus, CIRBP participates specifically in REM sleep homeostasis, whereas we do not find evidence for its participation in NREM sleep homeostatic mechanisms.

### OTHER MECHANISMS LINKING SLEEP-WAKE STATE TO CLOCK GENE EXPRESSION

Our results show that other pathways besides CIRBP must contribute to the sleep-wake driven changes in clock gene expression. Some suggestions for such pathways are shortly discussed below, as well considerations that could potentially account for the absence of a more widespread CIRBP dependent change in clock gene expression that we expected based on *in vitro* results.

*Rbm3* (RNA Binding Motif Protein 3), another cold-inducible transcript which is closely related to CIRBP, also conveys temperature information into high amplitude clock gene expression *in vitro* (Liu et al., 2013). Like *Cirbp*, its expression is sleep-wake driven (Wang et al., 2010). Thus, RBM3 might be another mechanism through which changes in sleep-wake state are linked to changes in clock gene expression and could therefore also explain the absence of a widespread CIRBP dependent SD-incurred change in clock gene expression. A follow-up study could address this possibility by quantifying the SD-evoked changes in clock gene expression in double *Cirbp*-*Rbm3* KO mice.

Heat shock factor 1 (*Hsf1*) is a member of the heat shock pathway and *in vitro* studies showed that it conveys temperature information to the circadian clock by initiating *Per2* transcription through binding to *Per2*’s upstream heat shock elements (Tamaru et al., 2011). Under undisturbed conditions, both *Hsf1* mRNA and protein levels are constitutively expressed, but the protein exhibits daily re-localization during the dark phase to the nucleus where it acts as a transcription factor (Reinke et al., 2008). Interestingly, CIRBP binds to the 3’UTR of *Hsf1* transcript [(Morf et al., 2012), see supplementary data therein], although it is unclear if this affects the transcriptional activity of HSF1. We found that SD induced a significant increase in *Hsf1* only in KO mice, which is congruent with the observation that the expression of two other transcripts downstream of HSF1, *Hsp90b* and *Hspa5/BiP*, was significantly amplified in KO mice after SD. Altogether, this suggests that the increased expression of *Per2* in KO mice might be linked to increased *Hsf1* expression, and underscores the presence of other temperature (and thus sleep-wake) driven pathways that can ultimately affect clock gene expression.

Beyond temperature, many other physiological changes occur during wakefulness that can subsequently affect clock gene expression. For example, oxygen consumption changes with sleep-wake state (Jung et al., 2011) and oxygen levels can modulate the expression of clock genes through HIF1α (Adamovich et al., 2017). Moreover, during SD, corticosterone levels increase which subsequently amplifies the expression of some, but not all, clock genes (Mongrain et al., 2010).

Another factor to consider is that the SD-incurred changes in clock gene expression depend also on an intact clock gene circuitry. For example, *Npas2* KO mice showed a reduced increase in *Per2* expression in the forebrain after SD (Franken et al., 2006), while *Cry1,2* double-KO mice display a larger increase in *Per2* expression after SD (Wisor et al., 2008). Thus, differences in clock gene circuitry, as suggested by the *in vitro* data (Liu et al., 2013, Morf et al., 2012), could also have contributed to the observed changes in clock gene expression after SD in *Cirbp* KO.

We could not corroborate the hepatic increase in heat shock transcripts (*Hsf1, Hsp90b* and *Hspa5*) and in *Per2* after SD as reported in other studies (Diessler et al., 2018, Maret et al., 2007), whereas we did confirm the SD-induced changes in *Cirbp, Rbm3-short, Dbp* and *Rev-erbα*. We cannot readily explain this lack of confirmation.

Finally, we would like to briefly address an obvious shortcoming. The hypothesis of this study is based on results obtained in a relatively simple biological model (*i.e.*, immortalized fibroblasts) and applied to a much more complex model (*i.e.*, cortices and livers of male mice). Unpublished observations on the circadian dynamics of the expression of CLOCK-BMAL1 target genes in the liver, such as *Rev-erbα* and *Dbp*, show an increased circadian amplitude in *Cirbp* KO mice; *i.e.* the opposite phenotype from that observed *in vitro* (Schibler et al., 2015). Along those lines, we could not consistently reproduce the importance of CIRBP in determining the *ext/com* ratio of *Sfpq* (see also Figure 4-B). Thus, *in vitro* findings will not always predict *in vivo* results, which could account for the lack of a widespread CIRBP-dependent change in clock gene expression after SD.

### CONCLUSION

This hypothesis-driven study explored whether the SD-induced changes in clock gene expression could be mediated through the cold-induced transcript CIRBP. After SD, the cortical expression of *Rev-erbα*, which we recently identified as a player in the sleep homeostat (Mang et al., 2016), was attenuated in *Cirbp* KO mice, whereas the expression of two other clock genes, *Per2* and *Clock*, was amplified. Thus, the SD induced changes in clock gene expression are, in part, modulated by CIRBP.

A large body of evidence has shown that clock genes are crucial for metabolism (reviewed in (Panda, 2016)). This is supported by the observation that disturbance of clock gene expression, through for example genetic manipulations in mice or shift work in humans, can lead to the development of metabolic disorders (Rudic et al., 2004) (Karlsson et al., 2001). Not only sleeping at the wrong time, but also sleeping too little or of poor quality can induce disturbed metabolic state both in rats (Barf et al., 2010) and humans (Copinschi et al., 2014). Because sleep loss affects clock gene expression (Franken, 2013), we propose that this could represent a common pathway through which both sleep and circadian disturbances lead to metabolic pathologies. It is thus of importance to determine the pathways through which a disturbed sleep-wake distribution affects clock gene expression. We show that temperature and CIRBP partake in this process, and we identified the expression of *Rev-erbα* as one of the genes affected by CIRBP. Genetic (Delezie et al., 2012) and pharmacological (Solt et al., 2012) studies have shown that this transcriptional repressor is important for healthy metabolic functioning. Further experiments could address the metabolic consequences of the attenuated response in *Rev-erbα* to sleep loss.

## Acknowledgements

We would like to thank Maxime Jan for his help in constructing the linear mixed model, our colleagues for their assistance with the sleep deprivations, Hannes Richter from the Genomic Technology Facility, for his support when setting up the RT-qPCR, David Gatfield and Bulak Arpat for fueling insightful discussions, and Jun Fujita (Kyoto University, Japan) for sharing the *Cirbp* KO mice. This study was performed at the University of Lausanne, Switzerland, and supported by the Swiss National Science Foundation (SNF n°146694 to PF supporting MMBH) and the state of Vaud (supporting MMBH, YE and PF).

## Material and Methods

### MICE AND HOUSING CONDITIONS

*Cirbp* KO mice, kindly provided by Prof Jun Fujita (Kyoto University, Japan), were maintained on a C57BL6/J background. In these mice, *Cirbp* exons were replaced by a TK-neo gene through homologous recombination in D3 embryonic stem cells, resulting in the absence of the *Cirbp* transcript and protein (Masuda et al., 2012). Breeding couples or trios consisted of heterozygous male and female mice. WT littermates were used as controls. Throughout all the experiments, mice were individually housed in polycarbonate cages (31×18×18 cm) with food and water *ad libitum* and exposed to a 12 h light/12 h dark cycle (70–90 lux). All experiments were approved by the Ethical Committee of the State of Vaud Veterinary Office Switzerland under license VD2743 and 3201.

### EEG/EMG AND THERMISTOR SURGERY

At the age of 9 to 13 weeks, 17 KO and 20 WT male mice were implanted with electroencephalogram (EEG) and electromyogram (EMG) electrodes (8 experimental cohorts). The surgery took place under deep xylazine/ ketamine anesthesia supplemented with isoflurane (1%) when necessary; for details see (Mang and Franken, 2012). Briefly, six gold-plated screws (diameter 1.1 mm) were screwed bilaterally into the skull over the frontal and parietal cortices. Two screws served as EEG electrodes and the remaining four anchored the electrode connector assembly to the skull. As EMG electrodes, two gold wires were inserted into the neck musculature. Of all EEG/EMG implanted mice, 8 KO and 9 WT mice were additionally implanted with a thermistor (serie P20AAA102M, General Electrics (currently Thermometrics), Northridge, California, USA) which was placed on top of the right cortex (2.5 mm lateral to the midline, 2.5 mm posterior to bregma). The EEG and EMG electrodes and thermistor were soldered to a connector and cemented to the skull. Mice recovered from surgery during 5–7 days before they were connected to the recording cables in their home cage for habituation, which was at least 6 days prior to the experiment. In total no less than 11 days were scheduled between surgery and start of experiment.

### EXPERIMENTAL PROTOCOL AND DATA ACQUISITION

EEG and EMG signals, Tcx and LMA were recorded continuously for 96 h. The recording started at light onset; i.e., Zeitgeber Time (ZT)0. During the first 48 h (days BL1 and BL2), mice were left undisturbed to establish a baseline. Starting at ZT0 of day 3, mice were sleep deprived by gentle handling for 6 hours (ZT0–6), as described in (Mang and Franken, 2012). The remaining 18 h of day 3 and the entire day 4 were considered as recovery (days REC1 and REC2, respectively). The analog EEG and EMG signals were amplified (2,000×), digitized at 2 kHz and subsequently down sampled to 200 Hz and stored. The EEG was subjected to a discrete Fourier transformation yielding power spectra (range: 0–100 Hz; frequency resolution: 0.25 Hz; time resolution: consecutive 4-sec epochs; window function: Hamming). Thermistors were supplied with a constant measuring current (*I*_*const*_ = 100 microA), and voltage (*V*) was measured at 10 Hz to calculate the median resistance (*Rt*) per 4-s epoch as in eq. (1).

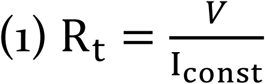

Each thermistor has an individual material constant, β. The resistance was measured at 25°C (*R*_*25*°*C*_) and 37°C (*R*_*37*°*C*_) by the manufacturer, and used to determine *β* as in eq.(2), with T values in Kelvin (°C + 273.15).

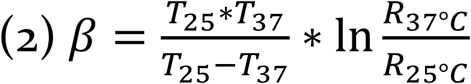

Following on eq. (2), the temperature (*t*) in °C can be calculated as described in eq. (3).

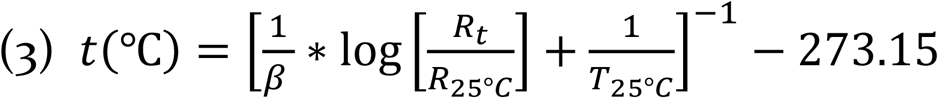

The EEG, EMG, and voltage across the thermistor were recorded with Hardware (EMBLA) and software (Somnologica-3) purchased from Medcare Flaga (EMBLA, ResMed, USA). LMA was detected with passive infrared sensors (Visonic Ltd, Tel Aviv, Israel) and quantified with ClockLab software (ClockLab, ActiMetrics, Wilmette, Illinois, USA).

### ANALYSIS OF LMA

To inspect the time course of LMA corrected for time-spent-awake, raw LMA was expressed per unit of waking in percentiles to which an equal amount of time-spent-awake contributed (as in Figure 3-G). The number of percentiles per recording period were chosen according to the prevalence of wakefulness, where 6 percentiles were used during the light phase and 12 during the dark phase, with the exception for 6 sections during the SD and 3 sections during the remaining 6hrs of the light phase of REC1. To assess genotype differences in LMA (Figure 6), the absolute number of movements were inspected. The LMA recordings of four mice (3 WT, 1 KO) were interrupted due to technical problems during the experiment, leaving data from 17 WT and 16 KO mice for analyses involving LMA.

We determined in 5 WT and 7 KO mice after the EEG-based sleep phenotyping their circadian period under at least two weeks of constant darkness. Period length was determined by Chi-squared test with ClockLab software (ClockLab, ActiMetrics, Wilmette, Illinois, USA).

### DETERMINATION OF BEHAVIORAL STATES

Offline, the mouse’s behavior was visually classified as ‘Wakefulness’, ‘REM sleep’, or ‘NREM sleep’ for consecutive 4-sec epochs based on the EEG and EMG signals, as previously described (Mang and Franken, 2012). Wakefulness was characterized by EEG activity of mixed frequency and low amplitude and variable muscle tone. NREM sleep was defined by synchronous activity in the delta frequency range (1–4 Hz), and low and stable muscle tone. REM sleep was characterized by regular theta oscillations (6–9 Hz) with low EMG activity. Waking was further differentiated into ‘quiet waking’ and ‘theta-dominated waking’ (TDW). TDW was determined based on the relative importance of power in the 6.5 to 12.0 Hz range to the overall power in the EEG of an artefact-free epoch scored as wakefulness, as described in (Vassalli and Franken, 2017). We refer to waking that is not classified as TDW as ‘quiet’ waking. Epochs containing EEG artefacts were marked according to the state in which they occurred and excluded from EEG spectral analysis but included in the sleep-wake distribution analyses. During the four day recording, 7.0 ± 0.9%, 2.1 ± 0.3% and 2.5 ± 0.2% of the epochs were scored as an artefact in waking, NREM, and REM sleep, respectively, and this did not differ between genotypes (t-tests, t(35)=1.77, p=0.09; t(35)=0.64, p=0.53; t(35)= 0.99, p=0.33, respectively).

### ANALYSIS OF CORTICAL TEMPERATURE

The raw T_cx_ data showed unexpected variation. Therefore, we inspected the inter-individual variation in daily amplitude and absolute T_cx_ levels. The latter was determined in two ways: i) by averaging T_cx_ during the last five hours of SD, thus minimizing the sleep-wake state incurred differences in T_cx_, and ii) by averaging T_cx_ during the 12h baseline light phase. These measures were highly correlated (R^2^=0.99; p<0.0001). Variation in the daily amplitude was quantified by averaging the difference between the highest and lowest hourly mean of T_cx_ of each of the two baseline days. No effect of genotype on absolute average T_cx_ or amplitude was detected (t-test, t(12)=0.61, p=0.55; t(12)=-0.63, p=0.54, respectively). Two mice (one of each genotype) exhibited a ca. 2-fold reduction in amplitude together with 2°C higher values during the SD relative to the other mice (Figure 7, pink symbols). Therefore, we excluded these two mice from subsequent T_cx_ analysis. Three other mice (2 WT and 1 KO) showed normal amplitude but overall lower absolute values (Figure 7, blue symbols). We corrected for this difference by raising their T_cx_ values by the difference between the T_cx_ reached in each of these 3 mice during the SD to the average T_cx_ reached over the same recording period in the remaining 9 mice. Of note, most of our T_cx_ analysis focuses on its relative sleep-wake dependent changes, which are not affected by differences in absolute T_cx_ values. Finally, the baseline T_cx_ data that was used to construct Figure 2B was based on 7 WT and 7 KO mice. During the recording, one KO mouse and one WT mouse had random fluctuations of T_cx_ beyond physiological reach and were therefore excluded from analysis involving the daily dynamics of T_cx_ (Figure 3). In the recovery, a KO mouse was excluded due to aberrant high T_cx_ that could not be accounted for by the sleep-wake distribution, leaving 6 WT and 5 KO mice for analyses involving REC1 and REC2.

**Figure 7.**
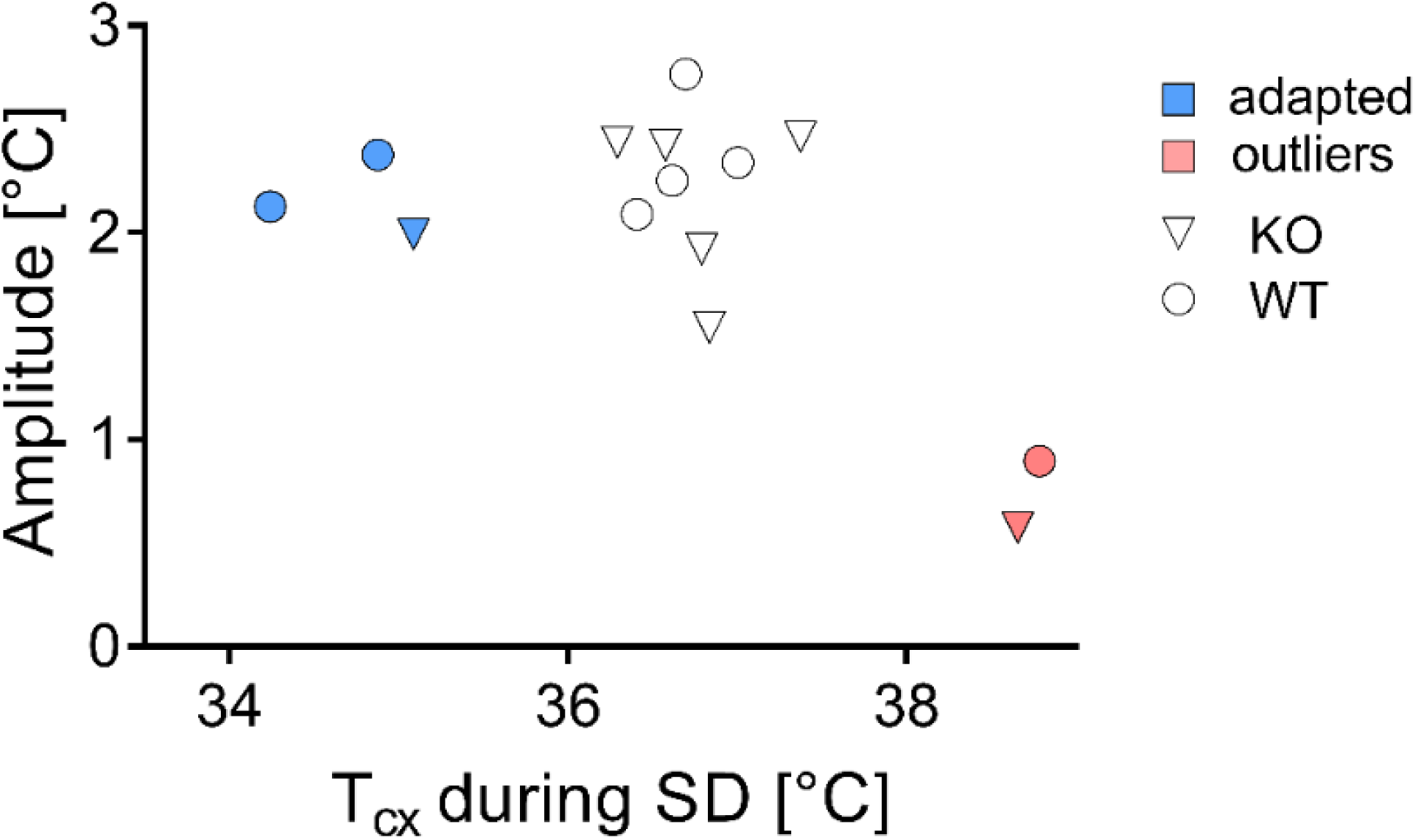
Assessment of amplitude and absolute values of cortical temperatures. Two outliers were detected (pink), whereas three others were corrected for their low values (blue).

Because visualization of all 4-s epochs occurring in 24-hour day is not compatible with the resolution of Figure 2-A, sleep-wake states were averaged per minute and assigned to either wake, NREM or REM sleep. Because REM sleep occurs less and in shorter bouts than waking and NREM sleep, this stage is slightly underrepresented in the hypnogram of Figure 2-A. T_cx_ was averaged per minute.

We inspected T_cx_ 1.5 min before and after sleep-wake transitions (*i.e.*, transitioning from wake to NREM sleep, NREM to REM sleep, REM sleep to wake and NREM sleep to wake). A sleep-wake transition was selected when the state before and after the transition lasted at least 8 epochs (*i.e.* >32 sec). With this criterion, an average of 38 wake to NREM sleep, 101 NREM sleep to REM sleep, 28 REM sleep to wake and 32 NREM sleep to wake transitions per mouse during the two baseline days was detected. The temperature profile of T_cx_ before and after the transition was constructed by depicting T_cx_ relative to the mean T_cx_ at a given sleep-wake transition (*i.e.*, the average of T_cx_ in the epoch before and after the sleep-wake transition). We subsequently constructed an individuals’ average change in T_cx_ for each sleep-wake transition. For this average, at least 10 traces were contributing at a given point in time to prevent skewing of the average individual temperature profile by few T_cx_ traces. Thus, the further from the sleep-wake state transition, the less epochs contributed to the average individual T_cx_ profile. One WT mouse exhibited an extreme drop in T_cx_ (−0.2°C in a 4-second epoch) after the transition from NREM sleep to wake in its average T_cx_ trace, but not in other sleep-wake transitions. We attributed this observation to a technical artefact and therefore this mouse was excluded from the NREM sleep to wake transitions.

The residuals of the correlation between waking and T_cx_ exhibited a circadian pattern under baseline conditions. We visualized the properties of this pattern further by fitting a sinewave through the data (Prism, non-linear regression; sine-wave with non-zero baseline; least squares fit).

### ANALYSIS OF EEG BASED ON BEHAVIORAL STATE

Unless otherwise stated, all mice (20 WT and 17 KO) were included in the analyses based on the EEG data. Spectral content of the EEG within sleep-wake states was calculated as follows. To account for inter-individual differences in overall EEG power, EEG spectra were expressed as a percentage of an individual reference value calculated as the total EEG power across 0.75-45 Hz and all sleep-wake states in the 48h baseline. This reference value was weighted so that for all mice the relative contribution of the three sleep-wake states (wake, NREM and REM sleep) to this reference value was equal.

Theta peak frequency (TPF) was calculated by determining the frequency at which power density peaks per 4-s epoch and subsequently averaged per individual. Power density peaks were quantified from 6.5 to 12.0 Hz band and from 5.5 to 12.0 Hz band for TDW and REM sleep, respectively.

Time course analysis of EEG delta power (*i.e.*, the mean EEG power density in the 0.75–4.0 Hz range in NREM sleep) during baseline and after SD was performed as described previously (Franken et al., 1999), and similar to the analysis of LMA per unit of waking. The light periods of BL1, BL2, and REC2 were divided into 12 percentiles, the REC1 light period (ZT6–12) into 8 sections, and all dark periods into 6 sections. The timing of these percentiles was based on the prevalence of NREM sleep. EEG delta power values in NREM sleep were averaged within each percentile and then expressed relative to the mean value reached in the last 4hr of the two main rest periods in baseline between ZT8–12. This reference was selected because delta power reaches lowest values at this time of the day and is least influenced by differences in prior history of sleep and wakefulness (see also (Franken et al., 1999)). In the time course of NREM delta power, one mouse (KO) demonstrated a strong decrease over the course of the experiment which could not be attributed to changes in the sleep-wake distribution. 9 out of the 12 delta power values during the light phase of REC2 in this mouse were outliers (MAD outlier test, consult (Leys et al., 2013) for details). This mouse was excluded from the analyses involving sleep homeostasis (Figure 5).

The effect of 6hr SD on subsequent time spent in NREM and REM sleep was assessed by calculating the recovery-baseline difference in sleep time per 1hr intervals.

### SIMULATING NREM EEG DELTA POWER [PROCESS S]

We applied a computational method to predict the change in delta power during NREM sleep based on the sleep-wake distribution as described before (Franken et al., 2001). Process S is exponentially increasing with time constant τ_*i*_ during waking and REM sleep, and exponentially decreasing by *τ*_*d*_ during NREM sleep (eq. (4) and (5), respectively).

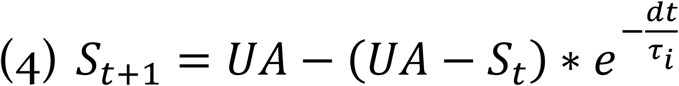

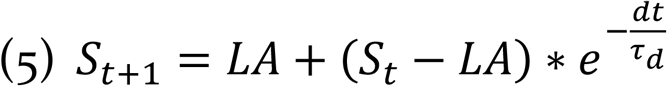

In these simulations, UA represents the upper asymptote, LA the lower asymptote and *dt* the time step of the iteration (4 seconds). Both asymptotes were estimated for each individual mouse. The upper asymptote was based on the 99% level of the relative frequency distribution of delta power reached in all 4s epochs scored as NREM sleep in the 4-day recording. As an estimate of the lower asymptote, the intersection of the distribution of delta power values in NREM sleep with REM sleep was taken. At the start of the simulation, an iteration through the first 24-hr (BL1) was performed with S_0_=150 at t=0. The value reached after 24-hrs is independent of S_0_ at t=0 and, assuming a steady state during baseline, reflects Process S at the start of the baseline for a given combination of time constants.

The fit was optimized by minimizing the mean squared difference of simulated and observed NREM delta power for a range of *T*_*i*_: 1-25 h, step size 0.125h; *T*_*d*_: 0.1-5.0 h, step size 0.025h; *i.e.* the simulation was run for all 38’021 combinations of *T*_*i*_ and *T*_*d*_ for each mouse. The combination of *T*_*i*_ and *T*_*d*_ giving the best fit was used to assess differences in process S between genotypes.

We noted a subtle but consistent linear discrepancy in the alignment of the simulated Process S to the measured NREM delta power values at the end of the light phase on BL1, BL2 and REC2 (Pearson correlation, slope≠0: 1 sample t-test; t(35)=-4.38, p=0.0001). This change correlated well with the day-to-day changes in total spectral power in the EEG calculated across all sleep-wake states in BL1, BL2, and REC2 (Pearson correlation: R^2^=0.70, p<0.0001; n=36). There was no effect of genotype on slope (Δ delta power %/h; students’ t-test; t(34)=0.62; p=0.54; WT:-0.086±0.027; KO:-0.065±0.021) or intercept (t(34)=-0.88; p=0.38; WT: 101.5±0.62; KO: 100.7±0.56; WT: n=20, KO: n=17). We attributed these linear changes to be of non-biological origin and detrended the measured NREM delta power values before optimizing the fit between observed and simulated delta power.

### GENE EXPRESSION IN LIVER AND BRAIN

Five mice of each genotype (n=15 per genotype in total) were sacrificed either prior to SD (ZT0), at ZT6 allowing them to sleep *ad lib* (*i.e.* without SD; ZT6-NSD), or at ZT6 after 6h SD (ZT6-SD) across four experimental cohorts. Mice were randomly assigned to one of the three experimental conditions. Genes of interest included transcripts affected by SD (Maret et al., 2007, Mongrain et al., 2010) and/or by the presence of CIRBP (Liu et al., 2013, Morf et al., 2012) with a special interest for clock genes. Specific forward and reverse primers and Taqman probes were designed (seeSupplementary File 1) to quantify mRNA.

Upon sacrifice, both the cerebral cortex and liver were extracted and immediately flash frozen in liquid nitrogen. Samples were stored at −80°C. RNA from cortex was extracted and purified using the RNeasy Lipid Tissue Mini Kit 50 (QIAGEN, Hombrechtikon, Switzerland); RNA from liver was extracted and purified using the RNeasy Plus Mini Kit 50 (QIAGEN, Hombrechtikon, Switzerland), according to manufacturer’s instructions. RNA quantity (NanoDrop ND-1000 spectrophotometer; Thermo Scientific, Wilmington, NC, USA) and integrity (Fragment Analyzer, Advanced Analytical, Ankeny, IA, USA) was measured and verified for each sample. 1000 ng of purified total RNA was reverse-transcribed in 20μL using a mix of First-strand buffer, DTT 0.1M, random primers 0.25μg/μl, dNTP 10mM, RNAzin Plus RNase Inhibitor and Superscript II reverse transcriptase (Invitrogen, Life Technologies, Zug, Switzerland) according to manufacturers’ procedures. The cDNA was diluted 10 times in Tris 10 mM pH 8.0, and 2μL of the diluted cDNA was amplified in a 10μL TaqMan reaction in technical triplicates on an ABI PRISM HT 7900 detection system (Applied Biosystems, Life Technologies, Zug, Switzerland). Cycler conditions were: 2 min at 50°C, 10 min at 95°C followed by 45 cycles at 95°C for 15 s and 60°C for 1 min. Standard curves were calculated to determine the amplification efficiency (E). A sample maximization strategy was used where all biological replicates of one tissue were amplified for two genes per plate. Gene expression levels were normalized to two reference genes (cortex: *Eef1a* and *Gapdh*: M=0.23, CV=0.09 and liver: *Gadph* and *Tbp;* M=0.32, CV=0.11) using QbasePLUS software (Biogazelle, Zwijnaarde, Belgium). *Rbm3* isoforms were in a separate run quantified in liver and cortex, again with their housekeeping genes (same as previously; cortex: M=0.22, and CV=0.08; liver: M=0.13, CV=0.05). Transcripts with an average Ct-value>30 were omitted from analysis (in KO and WT livers: *Rbm3, Dusp4, Homer1a* and *Npas2*; in cortex and liver of KO mice: *Cirbp*). Results are expressed as normalized relative quantity (NRQ) which based on the overall mean expression per gene, which was set at 1.0 (Hellemans et al., 2007).

CIRBP affects the poly-adenylation sites of several transcripts (Liu et al., 2013). We explored if this newly discovered role of CIRBP could be corroborated in our study by focusing on the transcript Splicing factor, proline and glutamine rich (*Sfpq*) which exhibits CIRBP-dependent alternative poly-adenylation (APA) ((Liu et al., 2013), see their Supplemental Fig4-5). We calculated the ratio of the prevalence of the external 3’UTR region over the common region according to eq. (6),

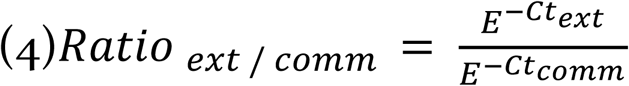

where E is the amplification efficiency and Ct_ext_ and Ct_comm_ the number of cycles for the detection of the extended and common isoform, respectively.

### STATISTICS

Statistics were performed in R (version 3.3.2) and Prism (version 7.0). The threshold of significance was set at p=0.05, and all statistics were solely performed on biological replicates. To be more specific, our RT-qPCR data stems from 5 biological replicates, whereas amplification of cDNA from one biological replicate is performed in three technical replicates. Deviations from the mean are representing standard error of the mean. The distribution of the LMA data was normalized by a log_2_ transformation on the hourly values, allowing for subsequent parametric analyses on the relationship between T_cx_ and LMA as in Figure 3. Time course data were analyzed by 1- and 2-way repeated measures (RM) analysis of variance (ANOVA) with as factors ‘time’ and ‘genotype’ (GT). Upon significance, post-hoc Fisher LSD tests were computed. Differences between BL and REC values within genotype were computed by paired t-tests. EEG spectra were also analyzed by 1- and 2-way RM ANOVA with as factors ‘time’ or ‘frequency’ and ‘GT’. When GT or its interaction with time or frequency reached significance, post-hoc t-tests were computed. The above-mentioned analyses were all performed in Prism. Missing values in the sleep-wake transition data of Figure 2-B led to exclusion of all data further away from the transition to perform a RM ANOVA.

Correlation coefficients of linear regression were calculated in Prism over all hourly values of LMA, T_cx_ and waking per genotype (96 per mice). To compare slopes of regression lines between genotypes, an ANCOVA was applied based on (Zar, 1984) and run in Prism. To quantify the contribution of waking and LMA independent from each other to T_cx_, a partial correlation was performed (R software; package ‘ppcor’, function pcor.test). Mixed model analysis was performed with factors LMA (log_2_ transformed), waking, and genotype (R packages ‘lme4’, ‘lmer’, ‘lmerTest’, and ‘MuMIn’). Model1 quantified the predictive power of waking, Model2 of waking and LMA per unit of waking (LMA/Waking) and Model3 of waking, LMA/Waking and its interaction, to predict T_cx_. Predictive power of models was compared with Chi-squared tests by assessing the statistical significance in the reduction of residual sum of squares between two models ordered by complexity; *i.e.* Model1 was compared to Model2, and upon significance, Model2 was compared to Model3. Goodness-of-fit was assessed by the marginal R-squared (R^2^_m_) which explains the effect of the fixed factors only, and the conditional R-squared (R^2^_c_), which considers the individual variance as well and is therefore more biological relevant. Hence, in the results section only the R^2^_c_ values are reported.

For the molecular data, the qPCR NRQ values were log_2_-transformed to normalize the distribution. Genotype differences at ZT0 were tested with a t-test. The effect of SD and genotype at ZT6 was assessed by 2-way ANOVA with post-hoc Fisher LSD tests upon significance. One outlier (WT, cortex) in the *ext/com* ratio analyses was detected by the Grubbs outliers test (α < 0.05) and excluded.

**Supplementary File 1**: sequences of the forward and reverse primer and probe used for the RT-qPCR

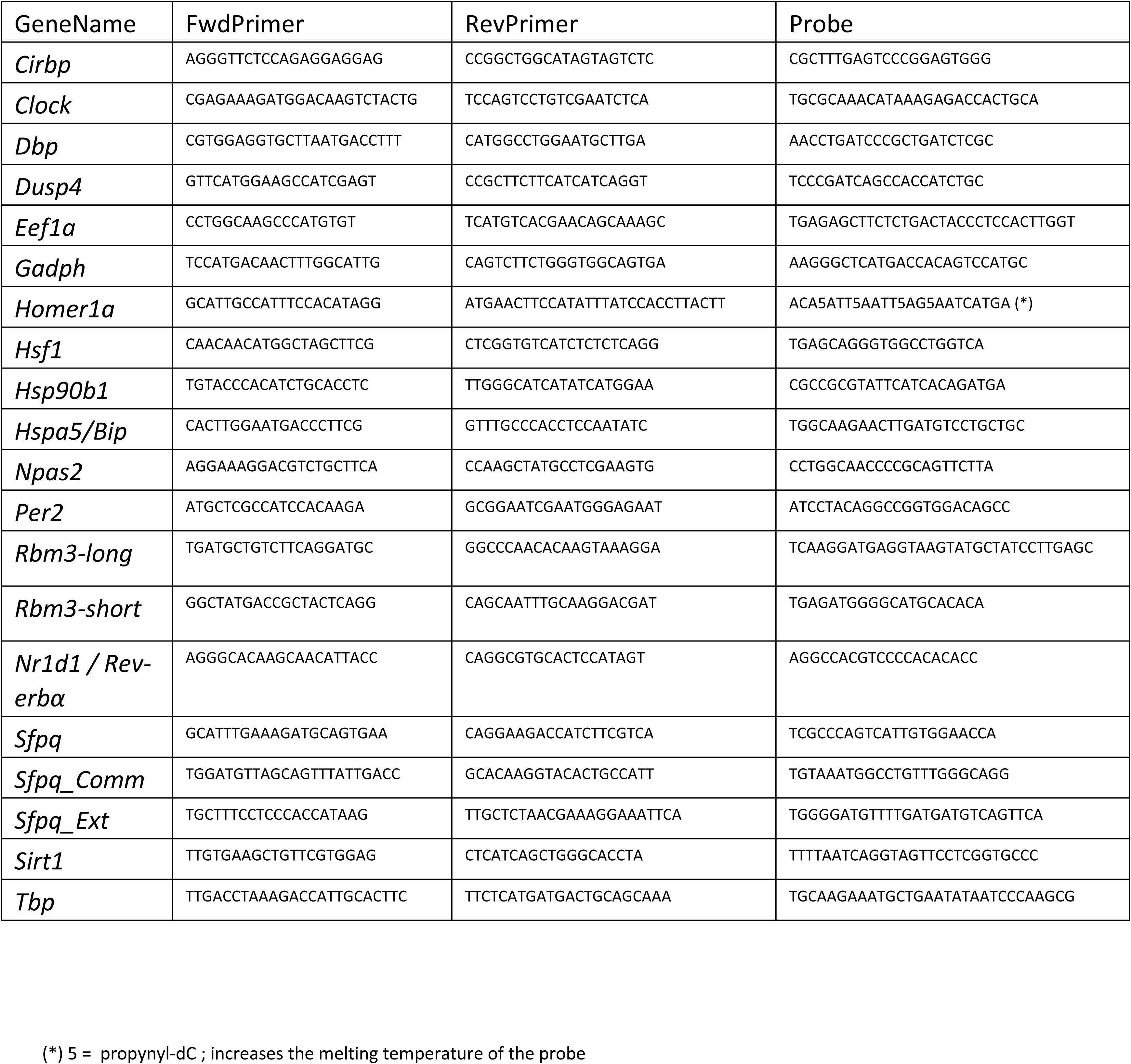

---

**Figure 3, supplement 1.**
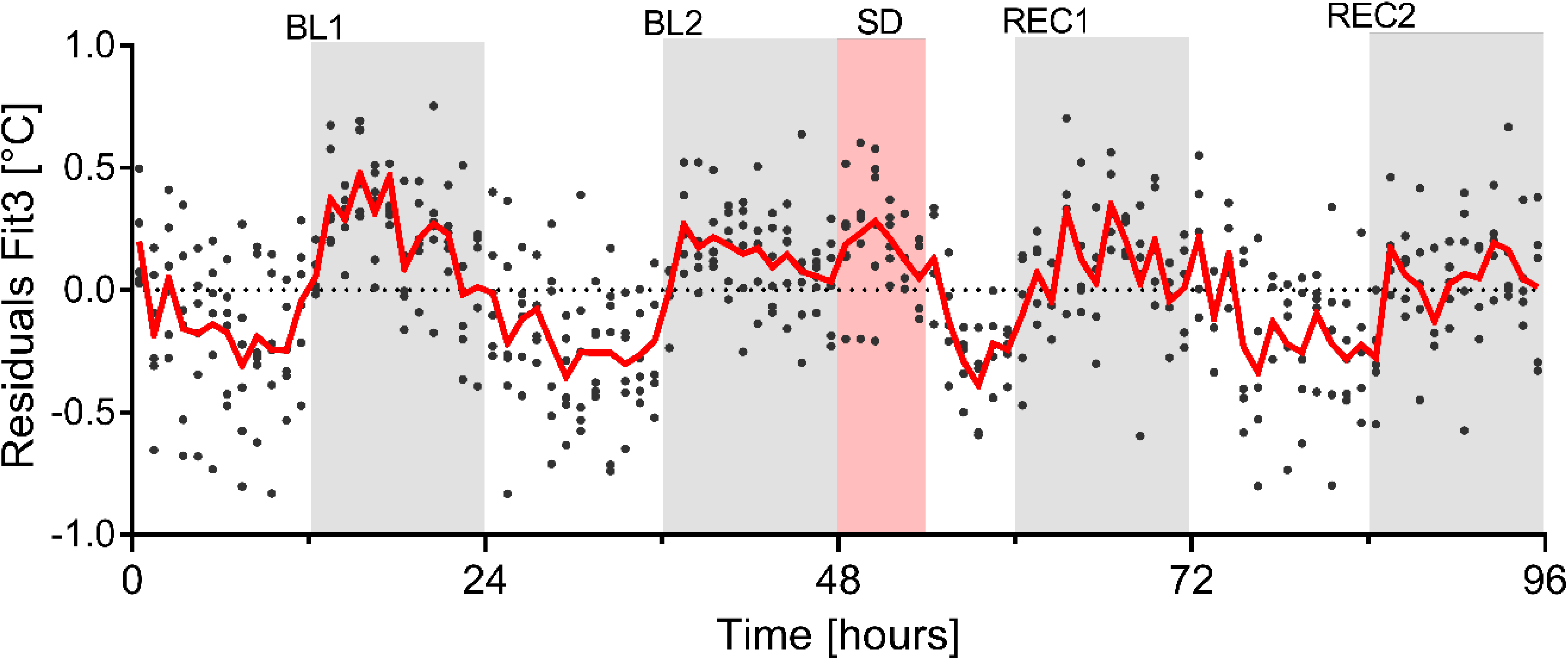
The residuals of the optimized mixed linear model. (Model3; see Results section) still show a pattern alike the residuals in Figure 3-F. Red line depicts the mean value determined every hour, each dot represents an hourly residual value per mouse.

**Figure 4, supplement 1.**
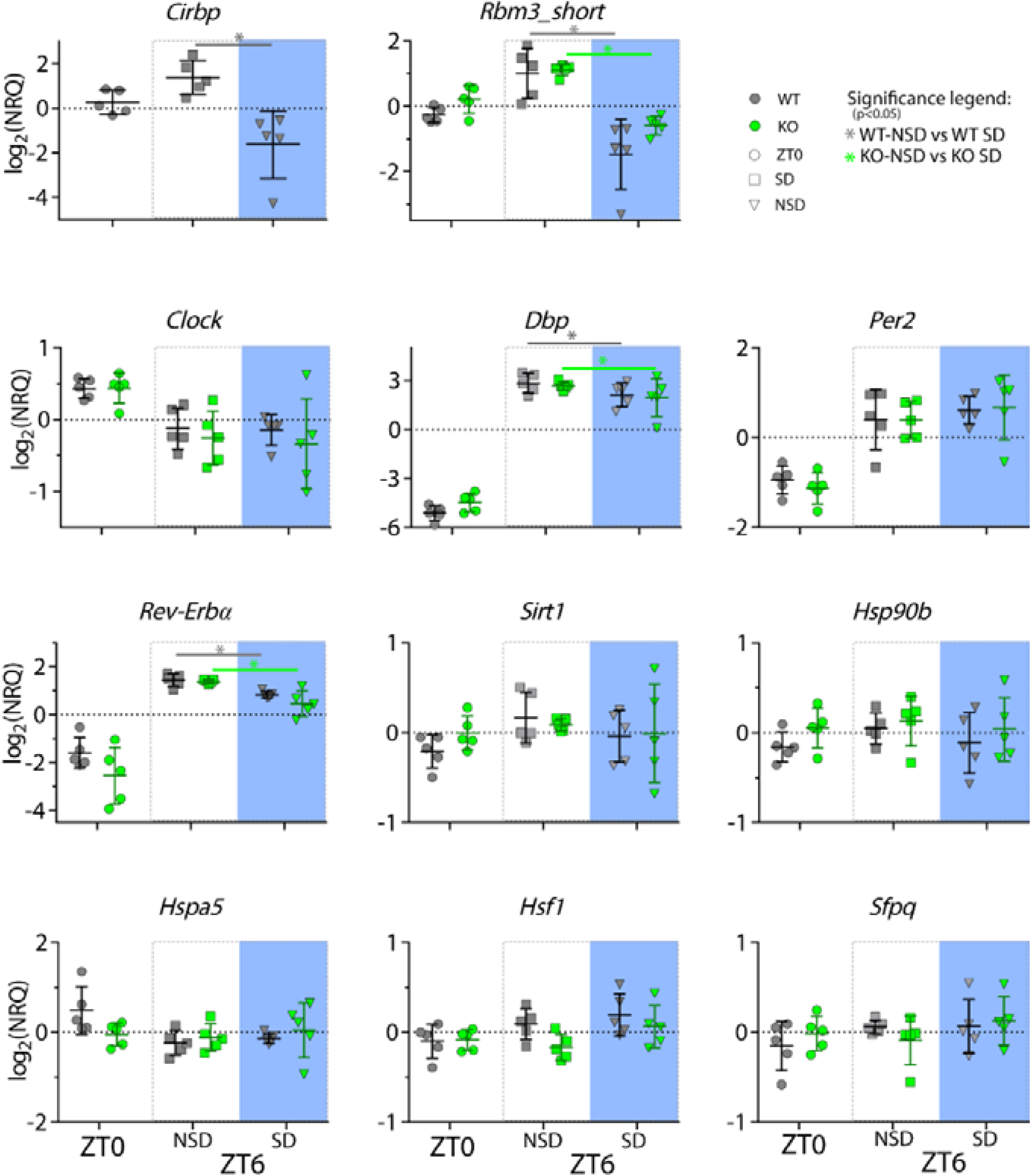
Changes in transcripts incurred by the absence of CIRBP and/or SD in the liver. Legend same as in Figure 4. See Table 1 for statistics.

**Figure 4, supplement 2.**
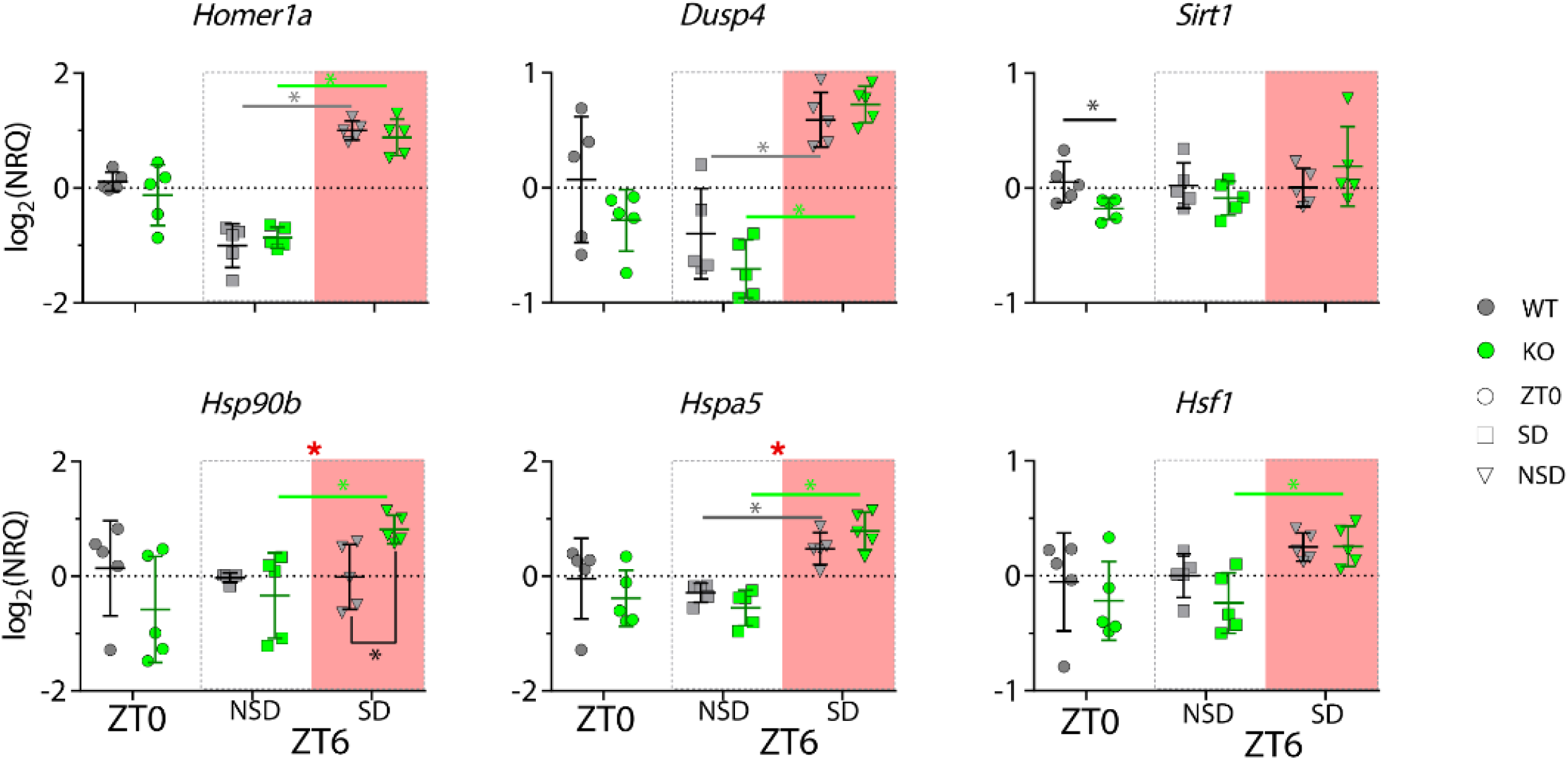
Changes in transcripts incurred by the absence of CIRBP and/or SD in the cortex. Legend same as in Figure 4. See Table 1 for statistics.

**Figure 6, supplement 1.**
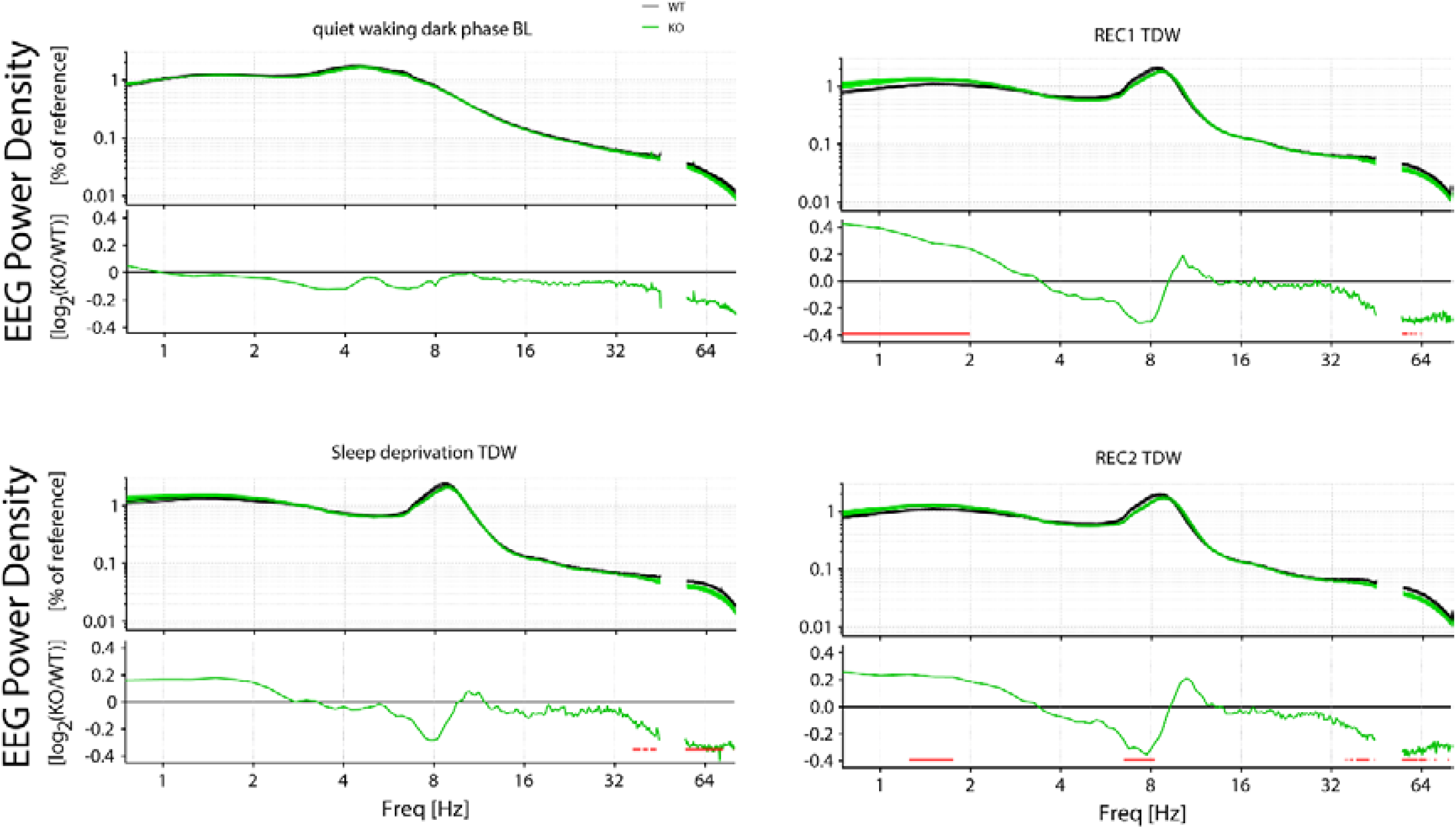
Changes in the EEG spectra are observed in TDW, but not in quiet waking. Spectral composition of *Cirbp* KO (green lines and areas; n=17) and WT (black line, grey areas; n=20) mice at different times of the experiment (indicated by the title, BL: baseline, REC1: recovery1, REC2: recovery2). In each graph, the top panel depicts the spectral composition whereas in the lower panel, the KO spectral composition relative to the WT is shown (ratio KO/WT). Red symbols in lower panel indicate significant difference for the frequency bins (post-hoc t-test, p<0.05). **<u>‘</u>**<u>Quiet’ waking baseline dark phase:</u> No effect of GT or an interaction effect (2-way RM ANOVA, GT*Freq: F(278, 9730)=0.81, P=0.99) on the spectral composition of quit waking in the dark phase of the baseline. <u>Sleep deprivation TDW</u>: significant interaction between GT and Freq (F (278, 9730) = 1.722, P<0.0001). <u>REC1 TDW:</u> significant interaction: F (278, 9730) = 2.984, P<0.0001; <u>REC2 TDW</u>: significant interaction F(278, 9730)=3.083, P<0.0001).

**Figure 6, supplement 2.**
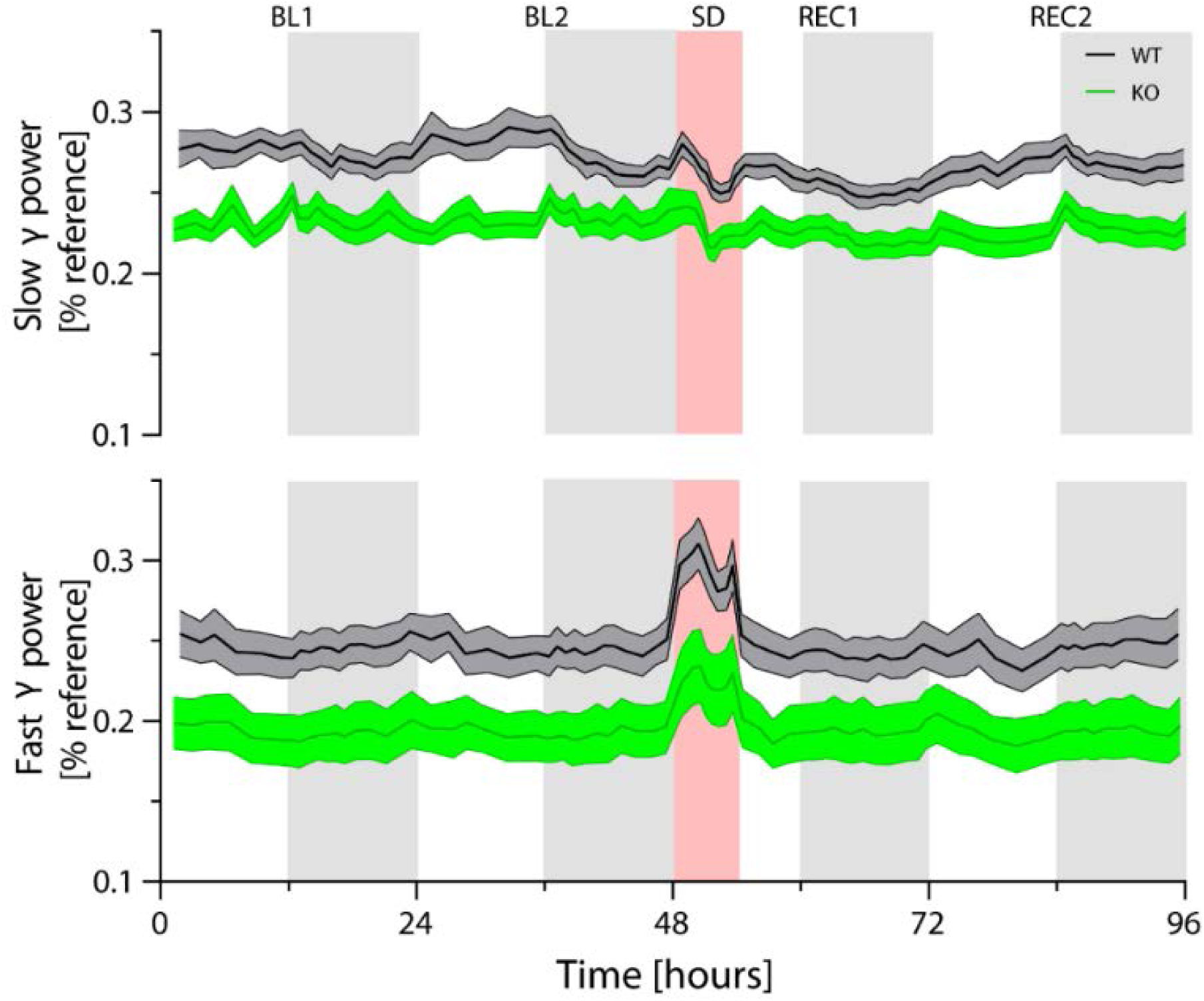
Slow and fast gamma power over the course of the experiment. *Cirbp* KO (green lines and areas) and WT (black line, grey areas) mice (areas span ±1SEM range). BL: baseline, SD: Sleep deprivation, REC: Recovery. Power in both slow [32-45Hz] and fast [55-80Hz] gamma is significantly reduced over the course of the experiment in *Cirbp* KO mice (2-way ANOVA: factor GT: slow gamma: F(1,34)=11.9, p=0.002; fast gamma: F(1,33)=6.6, p=0.01). Like the analysis of delta power, power in the gamma bands is calculated based on intervals to ensure an equal contribution of epochs to each time point: 6 intervals in the light phase, 12 in the dark periods, 8 in the sleep deprivation and 4 during the recovery light period.

